# The exchange of vitamin B_1_ and its biosynthesis intermediates in synthetic microbial communities shapes the community composition and reveals complexities of nutrient sharing

**DOI:** 10.1101/2021.09.29.462401

**Authors:** Rupali R. M. Sathe, Ryan W. Paerl, Amrita B. Hazra

## Abstract

Microbial communities occupy diverse niches in nature, and exchanges of metabolites such as carbon sources, amino acids, and vitamins occur routinely among the community members. While large-scale metagenomic and metabolomic studies shed some light on these exchanges, the contribution of individual species and the molecular details of specific interactions are difficult to track. Here, we explore the molecular picture of vitamin B_1_ (thiamin) metabolism occurring in synthetic communities of *Escherichia coli* thiamin auxotrophs which engage in the exchange of thiamin and its biosynthesis intermediates. In *E. coli*, the two parts of thiamin – the 4-amino-5-hydroxymethyl-2-methylpyrimidine and the 4-methyl-5-(2-hydroxyethyl)thiazole – are synthesized by separate pathways using enzymes ThiC and ThiG, respectively, and are then joined by ThiE to form thiamin. We observed that even though *E. coli ΔthiC, ΔthiE*, and *ΔthiG* mutants are thiamin auxotrophs, co-cultures of *ΔthiC*-*ΔthiE* and *ΔthiC*-*ΔthiG* grow in a thiamin-deficient minimal medium, whereas the *ΔthiE*-*ΔthiG* co-culture does not. Analysis of the exchange of thiamin and its intermediates in *Vibrio anguillarum* co-cultures, and in mixed co-cultures of *V. anguillarum* and *E. coli* revealed that the general pattern of thiamin metabolism and exchange among microbes is conserved across species. Specifically, the microorganisms exchange HMP and thiamin easily among themselves but not THZ. Furthermore, we observe that the availability of exogenous thiamin in the media affects whether these strains interact with each other or grow independently. This underscores the importance of the exchange of essential metabolites as a defining factor in building and modulating synthetic or natural microbial communities.

## Introduction

Microorganisms inhabit diverse natural habitats and ecosystems, and are engaged in a multitude of interactions, including sharing and competing for essential nutrients. Microbial communities or consortia are shaped via these positive and/ or negative interactions, and have their own unique metabolic network that is defined by the spatial distribution, physiology, and availability of nutrients among the microbial participants (1– 3). The exchange of biomolecules such as sugars, nucleobases, amino acids, vitamins, electron acceptors, fermentation byproducts, and metal-chelating siderophores are found to occur between members of natural and synthetic microbial consortia (1, 4–8). Some members of a microbial consortia may stop synthesizing a metabolite that is readily available in their environment, and eventually become auxotrophic for that nutrient (8, 9). Auxotrophy is beneficial for an individual organism as it allows for the reduction in metabolic burden and/ or genome size (10, 11). For example, in experiments with *Escherichia coli*, about 13% of mutants auxotrophic for vitamins, amino acids and nucleotides show a higher fitness than the wild-type strain when the missing nutrient is provided exogenously in sufficient quantities in the growth medium (12). Another study shows that the co-evolution of a co-culture of the sulfate-reducing bacterium *Desulfovibrio vulgaris* and an archaea *Methanococcus maripaludis* over 10^3^ generations leads to loss-of-function mutations in the sulfur-reducing genes in *D. vulgaris*. Further, deleting these genes shows an increased yield of the corresponding *D. vulgaris* strains when compared to the wild-type strain in the co-cultures (13). Conversely, microorganisms that produce a metabolite to share or exchange with their fellow community dwellers secure a position of prominence as they become indispensable for the consortium (14). *Ruminococcus bromii*, a starch-degrading bacterium associated with the human gut, converts starch to sugars that are substrates for other gut bacteria, thus playing the role of a keystone species (15). Thus, auxotrophy and metabolite sharing are important features for the formation and sustenance of microbial communities, and synthetic communities developed for biotechnological applications or for studying microbe-microbe interactions are often designed using these principles.

One of the commonly shared metabolites in microbial communities are vitamins (1, 16, 17). The activated forms of vitamins play an indispensable role as cofactors for numerous enzymes in primary metabolism across all domains of life. Several metagenomic analyses reveal that the water-soluble group B vitamins are readily exchanged in marine microbial communities, the human gut microbiota, and communities associated with insects and other hosts (17–20). Of these, vitamin B_1_ (also known as thiamin) is an important member of the group B vitamins that plays an essential role in carbohydrate, amino acid, and lipid metabolism by assisting enzymes in conducting “impossible” decarboxylations (21). In the human gut microbiome, B_1_ auxotrophy appears to be most widespread at both the genus and family level (17).

Even though most bacteria, plants and eukaryotes such as fungi are capable of thiamin biosynthesis, many other organisms are unable to produce it and instead acquire it from their surroundings or their diet. The structure of thiamin consists of a 5-membered 4-methyl-5-(2-hydroxyethyl)thiazole ring (THZ) and a 6-membered 4-amino-5-hydroxymethyl-2-methylpyrimidine ring (HMP) which are synthesized in the natural world in their phosphorylated forms, THZ-P and HMP-P, respectively, in two distinct branches of the thiamin biosynthesis pathway. The HMP-P is further phosphorylated to HMP-PP, following which the diphosphate is displaced by an attack via the THZ-P ring nitrogen to form a methylene bridge between the two rings to yield thiamin monophosphate (TMP) (Figure 1A). A final phosphorylation of TMP yields thiamin diphosphate (TDP), which is used as a cofactor by enzymes for cellular metabolism.

**Figure 1.**
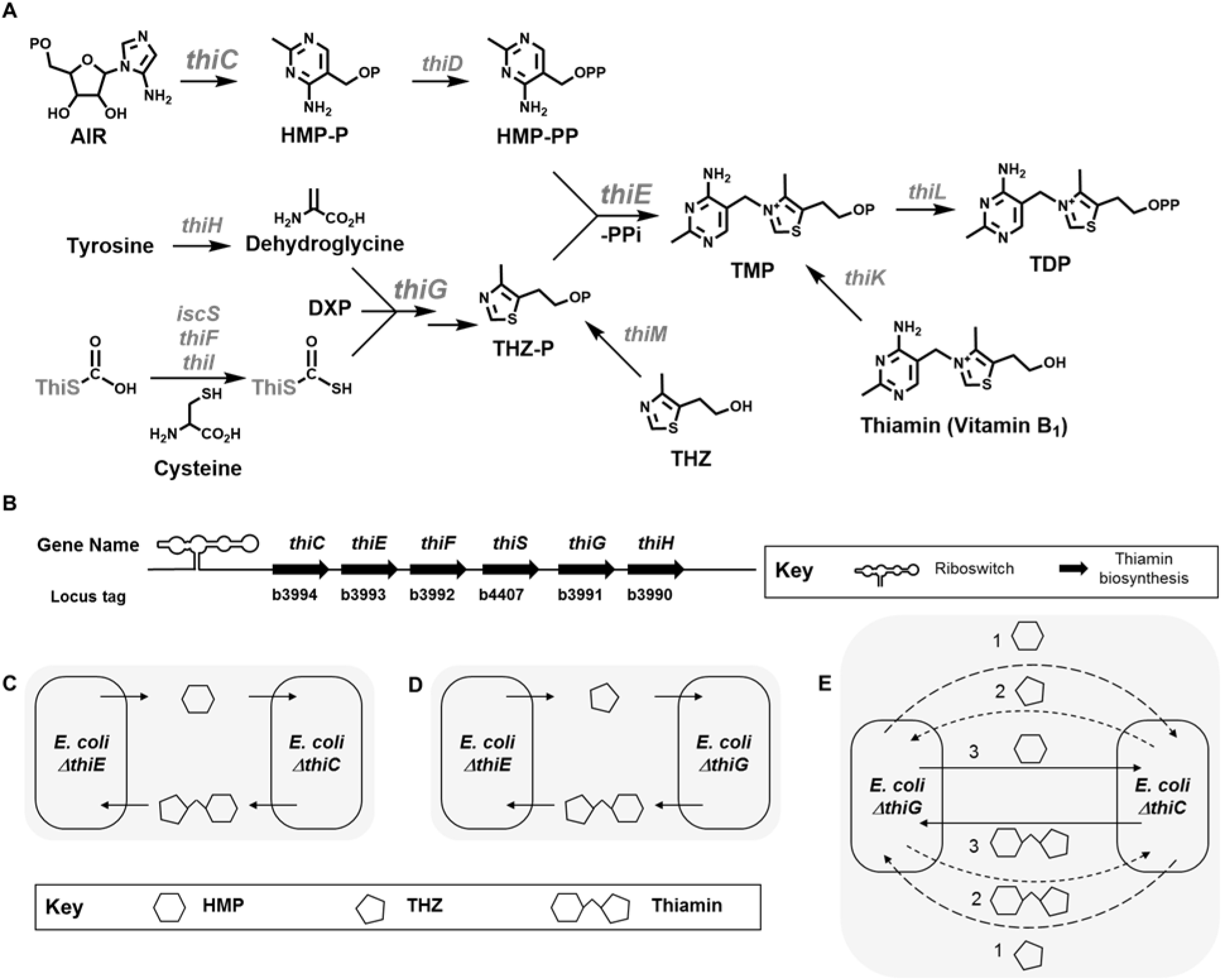
An overview of the thiamin biosynthesis pathway in *E. coli* str. K-12 substr. MG1655. **(A)** Major and relevant steps in the thiamin biosynthesis pathway are depicted. All phosphate groups are indicated as P. The 4-amino-5-hydroxy-2-methylpyrimidine phosphate (HMP-P) ring is formed by rearrangement of its precursor aminoimidazole ribotide (AIR), by the enzyme ThiC. The 4-methyl-5-(2-hydroxyethyl)thiazole ring (THZ) is formed by the enzyme ThiG. The sulfur in the THZ ring is transferred through a cascade of enzymes IscS, ThiI, ThiF and ThiS in the form of a thiocarboxylate moiety. The 1-deoxyxylulose-5-phosphate is synthesized by Dxs, and the ThiH enzyme converts tyrosine to dehydroglycine. ThiD, ThiM, and ThiK act as kinases for HMP-P, THZ, and thiamin respectively. The enzyme ThiE attaches the HMP-PP and the THZ-P rings together in the final steps of the pathway to form thiamin monophosphate (TMP) and ThiL phosphorylates it to form the active form of the cofactor, thiamin pyrophosphate/ diphosphate (TDP). Abbreviations: AIR = 5’-phosphoribosyl-5-aminoimidazole, DXP = 1-deoxyxylulose-5-phosphate, HMP-P = 4-amino-5-hydroxymethyl-2-methylpyrimidine phosphate, HMP-PP = 4-amino-5-hydroxymethyl-2-methylpyrimidine diphosphate, THZ-P = 4-methyl-5-(2-hydroxyethyl)thiazole phosphate, THZ = 4-methyl-5-(2-hydroxyethyl)thiazole. Enzyme names are written in grey. **(B)** The arrangement of the genes involved in the *de novo* biosynthesis pathway for thiamin in *E. coli* K-12 MG1655. We hypothesize that **(C)** the co-culture of *EcΔthiC*-*EcΔthiE* strains can survive by exchanging HMP and thiamin, **(D)** the co-culture of *EcΔthiE*-*EcΔthiG* strains can survive by exchanging THZ and thiamin, and **(E)** the co-culture of numbers *EcΔthiC*-*EcΔthiG* strains can survive in three possible scenarios where metabolites are exchanged in following pairs: 1 – exchange of HMP and THZ, 2 – exchange of THZ and thiamin, and 3 – exchange of HMP and thiamin. Hexagon – HMP, pentagon – THZ, linked hexagon and pentagon – thiamin.

Thiamin and its intermediates THZ and HMP are stable under physiological conditions and are salvaged from the environment by organisms for producing thiamin. Metagenomic analysis of the human gut microbiome reveals that the thiamin biosynthesis and salvage pathways display the largest variety of intermediates and non-canonical metabolic precursors (17). Recent findings implicate HMP as an important metabolite in shaping marine algal and bacterial consortia (22). Additionally, studies show that there exist thiamin auxotrophs that lack thiamin transporters but instead contain putative transporters for the uptake of HMP and/ or THZ, which permit the salvage of these intermediates to produce thiamin (16, 23, 24). Examples of HMP and thiamin transporters and their uptake have been reported widely in literature (25–28). On the other hand, information on THZ uptake and exchange is limited to only a handful of studies that predict a THZ transporter and show the uptake of the precursor carboxythiazole (16, 25, 26, 29, 30).

The modular nature of thiamin biosynthesis where HMP and THZ are found to be independently synthesized and salvaged makes this pathway a unique candidate for studying metabolic crosstalk within microbial co-cultures. To experimentally validate some of these findings, we require a simple model system whose members are engaged in thiamin, THZ, and HMP exchange. Such a system will allow us to (i) understand thiamin biosynthesis at a community level, that is, beyond what occurs in individual organisms and (ii) establish the design principles of building synthetic communities sustained by thiamin biosynthesis and uptake with diverse biotechnological applications.

In this study, we create a series of thiamin-dependent synthetic co-cultures using *E. coli*, a gram-negative bacterium that is capable of *de novo* thiamin synthesis and salvage, and is a member of several environmental and enteric microbial communities. In *E. coli*, the formation of the HMP-P ring is catalyzed by the enzyme ThiC, the THZ-P ring is synthesized by a host of enzymes including ThiG, and subsequently, these rings are coupled together by ThiE to form TMP. Also, no known transporters and salvage enzymes of the HMP or THZ or their analogues are found in this organism, and only one known transporter ThiBPQ exists to facilitate thiamin transport (22, 25, 26, 30–32). We generated *E. coli* str. K-12 substr. MG1655 thiamin biosynthesis mutants - *ΔthiC, ΔthiE* and *ΔthiG -* which are impaired in *de novo* thiamin biosynthesis, and thus are thiamin auxotrophs. We then set up pairwise synthetic co-cultures of these three *E. coli* mutants to study their growth over short time periods. We also analyzed the exchange of thiamin, THZ, and HMP at a molecular level, and its effect on the community composition. Further, we studied similar co-cultures of another gammaproteobacterium *Vibrio anguillarum* and finally, mixed co-cultures of *E. coli* and *V. anguillarum* to understand the extent to which our findings on thiamin metabolism within the *E. coli* communities hold true for other bacteria.

A unique property of the thiamin-based synthetic consortia we have devised is that these are reliant on the exchange of precursors and intermediates within a single metabolic pathway, as compared to other synthetic co-culture studies in literature which involve exchange of molecules derived from two or more metabolic pathways (4, 6, 33). The advantages of a co-culture system which is based on the biosynthesis of a single metabolite are: (a) the growth conditions of individual strains are similar as they are auxotrophs for the same metabolite, (b) the regulation of the biosynthesis and uptake of individual intermediates along the pathway can be studied, and (c) coupling the results we observe from our system with genetic data from isolates and metagenomes has the potential to improve predictions of B1-related auxotrophy and metabolite exchange in natural systems.

Our results indicate that the rules of exchange of thiamin and its intermediates are broadly similar across organisms, and variations may be predicted based on growth conditions and the genome sequences of the interacting species. We also observe temporal changes in the ratios of the *thi*^*-*^ mutants in our co-cultures based on the ability of the strains to either make B_1_ or a B_1_ biosynthesis intermediate, or use exogenously added B_1_. Our findings inform on the physiology of single microbial members with regard to thiamin metabolism within the context of a microbial community. Finally, our study highlights the nature of interdependencies that arise from relying on acquiring essential metabolites from the environment or from fellow community members.

## Results

### *E. coli* thiamin biosynthesis auxotrophic mutants show concentration-dependent increase in growth when supplemented with thiamin or its biosynthesis intermediates

*E. coli* is capable of producing TDP *de novo* and also contains genes to salvage thiamin from its environment (Figure 1A). All the major genes for thiamin synthesis and salvage are found in three operons: (i) *thiCEFSGH*, which conducts *de novo* thiamin biosynthesis (Figure 1B), (ii) *thiMD*, which codes for kinases in the salvage pathway, and (iii) *thiBPQ*, which codes for an ABC-type thiamin transporter (34) (Figure S1A). All three operons are regulated by TDP-dependent riboswitches (35).

To begin our studies, we created three knockout *E. coli* K-12 MG1655 strains - *EcΔthiC, EcΔthiE* and *EcΔthiG* (referred to as the *thi*^*-*^ mutant strains) - and noted that all three strains grew without any growth disadvantage in a nutrient-rich medium (Figure S2A). Next, we tested their growth in a thiamin-deficient minimal medium containing M9 salts with glucose and NH_4_Cl as the carbon and nitrogen sources, respectively. We expected that they would require exogenously added thiamin for growth in this media, but to our surprise, all three strains survived well in the first passage (P1) from the nutrient-rich to the minimal medium (Figure S2C). A second passage (P2) of the mutants in the thiamin-deficient minimal medium showed significantly lesser growth as compared to the wild-type strain, and the third passage (P3) showed no growth, indicating that the *thi*^*-*^ mutant strains were indeed thiamin auxotrophs (Figure 2A-D, no thiamin added trace, Figure S2E). Our results match similar observations in literature, which note that thiamin stored inside the cells during their growth in rich media is carried over into a few generations of cell growth (36, 37). For all future experiments, the P2 cells were used as this allowed for us to have some cells from the controls for thiamin quantitation experiments while yet showing a sufficient difference in optical density (OD_600_) between the single culture and co-culture growth experiments.

**Figure 2.**
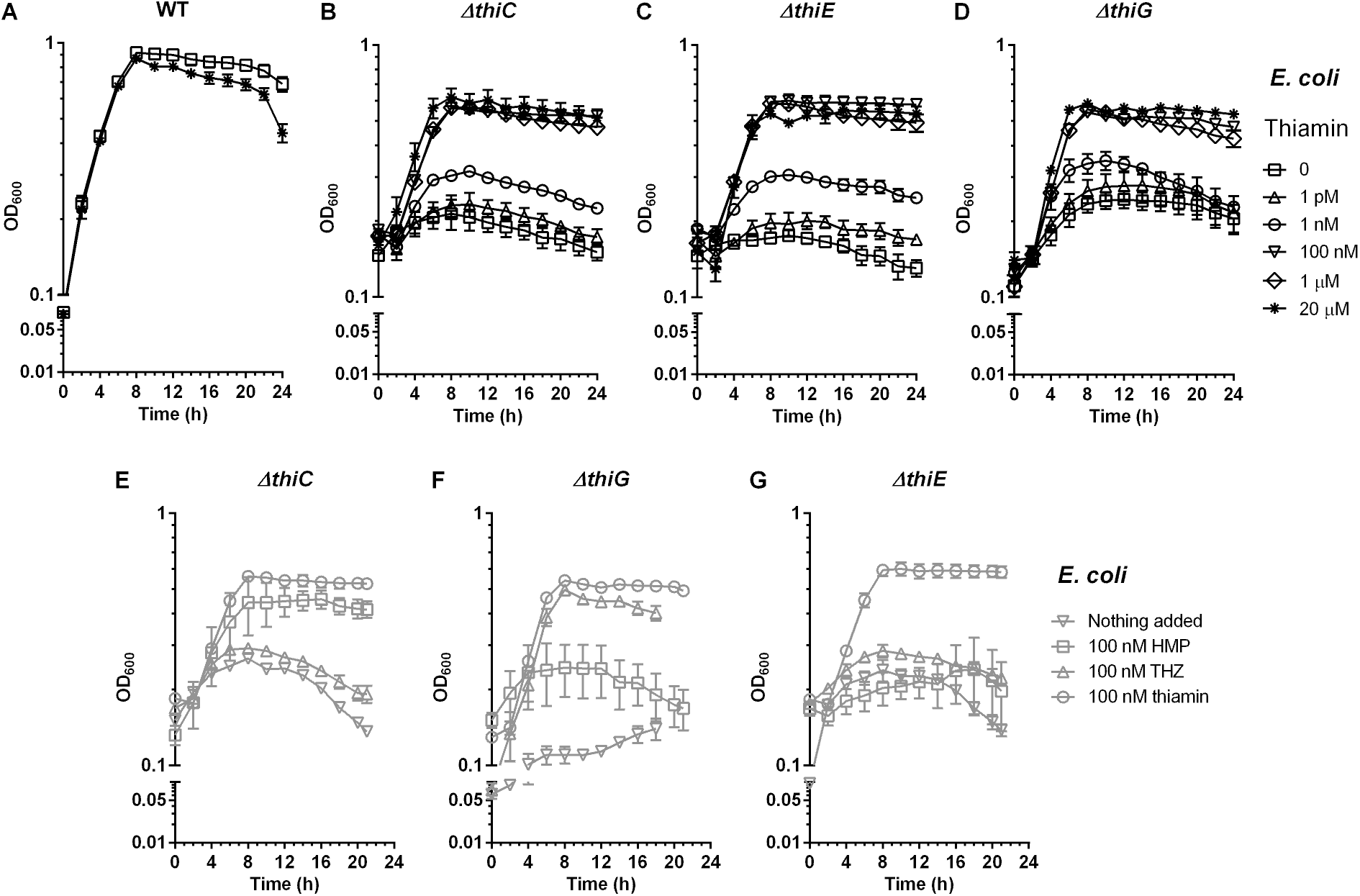
Supplementation of thiamin, HMP and THZ to the thiamin mutants of *E. coli* K-12 MG1655 in M9 medium. Growth phenotype of **(A)** the wild-type strain, **(B)** the *EcΔthiC* strain, **(C)** the *EcΔthiE* strain, and **(D)** the *EcΔthiG* strain. Symbols in panels **(A-D)** depict the following concentrations of thiamin: □ = Nothing added, ○ = 1 pM, Δ = 1 nM, ∇ = 100 nM, ◊ = 1 µM, * = 20 µM. Supplementation to **(E)** the *EcΔthiC* mutant, **(F)** the *EcΔthiG* mutant, and **(G)** the *EcΔthiE* mutant. Symbols in panels **(E-G)** depict the following: ∇ = no HMP/ THZ, □ = 100 nM HMP, Δ = 100 nM THZ, ○ = 100 nM thiamin. Means ± standard errors of the means from three independent experiments are plotted.

Next, to determine the minimum thiamin concentration required by the *thi*^*-*^ mutant strains, we tested their growth in minimal medium supplemented with thiamin concentrations ranging from 0 to 20 µM. We found that while these strains show low growth with up to 1 nM thiamin, they are able to achieve an OD_600_ of ∼0.6 with 100 nM, 1 µM and 20 µM thiamin (Figures 2B, 2C and 2D). This shows that thiamin is the growth-limiting nutrient for the *thi-* mutants. To ensure that thiamin is not limiting in our assays, all further experiments were conducted with 20 µM thiamin unless otherwise stated. We further complemented each knockout strain with a plasmid containing the deleted gene and confirmed that growth can be restored in these strains in minimal medium in the absence of thiamin, as also observed in previous literature studies (Figure S3) (38–40).

We expect the *EcΔthiC* and *EcΔthiG* strains to be impaired in the biosynthesis of the intermediates HMP and THZ, respectively, while the *EcΔthiE* mutant should not be able to link them together to synthesize thiamin. To test this, we fed HMP and THZ to the *thi-* mutants in varying concentrations ranging from 0-1 μM. The *EcΔthiC* strain survived only when supplemented with HMP but not with THZ (Figures 2E, S4), the *EcΔthiG* strain when supplemented with THZ but not with HMP (Figures 2F, S4), and the *EcΔthiE* strain was unable to survive with either HMP or THZ alone (Figures 2G, S4).This confirms that the metabolic phenotypes of the *thi-* mutants are correlated to their genotypes.

### Specific co-cultures of the thiamin biosynthesis mutants grow in minimal medium with no exogenously added thiamin

Next, we constructed three pairwise co-cultures *EcΔthiC*-*EcΔthiE* (*Ec-CE), EcΔthiC*-*EcΔthiG* (*Ec-CG)*, and *EcΔthiE*-*EcΔthiG* (*Ec-EG)* and studied their growth in thiamin-deficient minimal medium. We hypothesized that, if the *thi-* mutant strains can share thiamin biosynthesis intermediates among themselves and produce thiamin, the co-cultures will survive as opposed to the single cultures which are auxotrophic and perish under similar growth conditions. We started these co-cultures with 9:1, 1:1 and 1:9 ratios and observed their growth over a period of 24 hrs. We observed that the 1:9 ratio of the *Ec-CE* co-culture and the *Ec-CG* co-culture showed increased survival as compared to the individual pure cultures (Figure 3A and B, data for the 9:1 and 1:1 not shown). On the other hand, the *Ec-EG* co-culture showed no difference in growth when compared to their individual pure cultures (Figure 3C).

**Figure 3.**
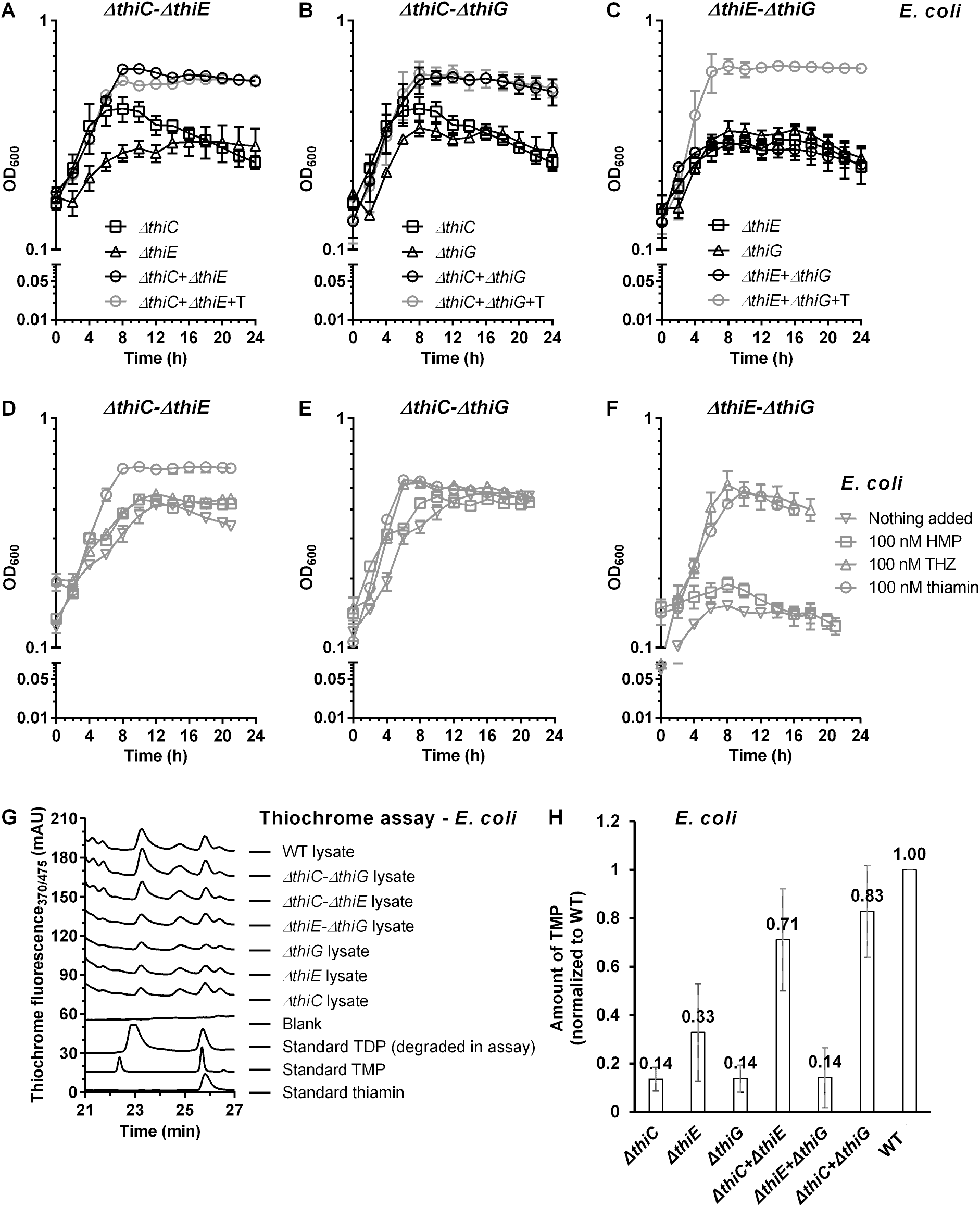
Thiamin biosynthesis mutants of *E. coli* K-12 MG1655 grow in pairwise co-cultures in thiamin-deficient M9 medium. Co-culture of **(A)** the *EcΔthiC*-*EcΔthiE* strains, **(B)** the *EcΔthiE*-*EcΔthiG* strains, and **(C)** the *EcΔthiC*-*EcΔthiG* strains. Supplementation to **(D)** the *ΔthiC-ΔthiE* co-culture, **(E)** the *ΔthiE-ΔthiG* co-culture, and **(F)** the *ΔthiC-ΔthiG* co-culture. Symbols in panels **(D-F)** depict the following: ∇ = no HMP/ THZ, □ = 100 nM HMP, Δ = 100 nM THZ, ○ = 100 nM thiamin. **(G)** HPLC of thiochrome assay samples to detect thiamin from co-culture lysates. **(H)** Amount of thiochrome monophosphate (TMP) normalized to that in the WT strain, detected in lysates of monocultures or co-cultures of the thiamin biosynthesis mutants grown in P2. Means ± standard errors of the means from three independent experiments are plotted.

There are several possibilities of exchange of thiamin and its intermediates that account for the survival of the *Ec-CE* and *Ec-CG* co-cultures (Figures 1C-E). The *EcΔthiC* strain cannot synthesize HMP, but if it can acquire it from its environment, it can combine the HMP with the THZ it synthesizes to form thiamin. Alternately, it can acquire thiamin directly from its environment. Similarly, the *EcΔthiG* strain cannot synthesize THZ, but it can grow if it acquires THZ or thiamin from its surrounding. On the other hand, the *EcΔthiE* strain can synthesize both the HMP and the THZ intermediates, but is unable to combine them to form thiamin and needs to acquire it from its growth medium. The growth observed in the *Ec-CE* co-culture can be explained only if the *EcΔthiE* strain supplemented the *EcΔthiC* strain with HMP, and the *EcΔthiC* strain in return supplemented the *EcΔthiE* strain with thiamin (Figure 1C, 3A). This indicates that both HMP and thiamin are likely being exchanged in the medium. On similar lines, the *Ec-EG* co-culture can grow only if the *EcΔthiE* strain supplemented the *EcΔthiG* strain with THZ, and the *EcΔthiG* strain in return, supplemented the *EcΔthiE* strain with thiamin (Figure 1D, 3C). As the *Ec-EG* co-culture does not grow, and we know that THZ and thiamin are salvaged by the *E. coli* cells based on our feeding studies and thiamin is also exchanged as per the results of the *Ec-CE* co-culture, this result suggests that THZ is possibly not present at a large enough concentration to be taken up by the *EcΔthiG* mutant. The absence of any annotated THZ transporters in *E. coli* also supports this hypothesis. Interestingly, the *Ec-CG* co-culture shows a distinct increase in its growth as compared to the individual pure cultures grown in thiamin-deficient medium (Figure 3B). The *Ec-CG* co-culture can grow in three scenarios: (i) the *EcΔthiC* strain and the *EcΔthiG* strain provided the other with THZ and HMP, respectively, or (ii) the *EcΔthiC* strain provided the *EcΔthiG* strain with THZ and, the *EcΔthiG* strain synthesized thiamin and provided it back to the *EcΔthiC*, or (iii) the *EcΔthiG* strain provided the *EcΔthiC* strain with HMP and, the *EcΔthiC* strain synthesized thiamin and provided it back to the *EcΔthiG* strain (Figure 1E).

As the data of the *Ec-EG* co-culture indicates that THZ is likely not being exchanged, only the third possibility remains for the *Ec-CG* co-culture, that is, HMP and thiamin are exchanged among the thiamin biosynthesis mutants. Incidentally, several reports in literature note the exchange or release of HMP among microbial communities, confirming our observation (23, 41).

Next, we compared between carbon sources to understand whether these results hold true across different growth conditions. In addition to glucose, we chose pyruvate and succinate as their utilization requires thiamin. Similar to what we observed with glucose, the *Ec-CE* and *Ec-CG* co-cultures showed growth in pyruvate and succinate minimal media without thiamin while the *Ec-EG* did not, and the growth of the *Ec-CG* co-culture was highest among the three (Figure S5). As the metabolism of glucose, pyruvate and succinate require thiamin-utilizing enzymes, our growth studies imply that the *Ec-CE* and *Ec-CG* co-cultures are able to synthesize thiamin. The *Ec-CE* co-culture showed lower growth in the presence of pyruvate and succinate as compared to glucose, and thus we continued with glucose as the carbon source for all further experiments.

### Analysis of the co-cultures demonstrate that the exchange of HMP and thiamin aids their survival

To probe the growth patterns observed for the *Ec-CE, Ec-CG* and *Ec-EG* co-cultures, we conducted a supplementation study with a range of HMP and THZ concentrations (Figures 3D-F, Figure S4). We observed that while the *Ec-CE* and *Ec-CG* co-cultures each show growth without or with supplementation with both molecules, the *Ec-EG* co-culture survives only when fed with THZ, but not with HMP (Figure 3F). Interestingly, the *EcΔthiG* mutant can survive with 1 nM THZ, whereas the *EcΔthiC* mutant requires 100 nM HMP to survive (Figures S4A, I, K). This indicates that *E. coli* differs in its ability to either acquire and/ or utilize the thiamin biosynthesis intermediates THZ and HMP. This result also sheds light on one of our preliminary observations that when the *Ec-CE* and *Ec-CG* co-cultures were started at a total OD_600_ of 0.01 in the P2 passage instead of 0.1, they were unable to survive (data not shown). We attribute this to the lack of an adequate pool of thiamin intermediates at the start that would allow the co-culture strains to begin dividing and cooperating, thus ensuring their survival.

### *De novo* biosynthesis of thiamin occurs within co-cultures

To verify that the growth of the co-cultures is due to the *de novo* biosynthesis of thiamin, we analyzed the lysates of the cells grown in thiamin-deficient media from the second passage P2 for single cultures and co-cultures for the presence of thiamin and its phosphorylated versions TMP and TDP. To do so, we used the thiochrome assay which employs an oxidation reaction under alkaline conditions to generate a fluorescent derivative of thiamin (42). Firstly, we noted that under the thiochrome assay conditions we used, the standard thiochrome diphosphate formed is unstable, and undergoes dephosphorylation (Figure 3G, HPLC trace). Next, we analyzed the lysates of the *Ec-CE* co-cultures and the *Ec-CG* co-cultures and noted that the levels of thiochrome monophosphate in them were significantly higher than their respective single cultures when measured at 24 hr, and similar to those in the wild-type *E. coli* cell lysate (Figure 3G, H). In contrast, the amounts of thiochrome monophosphate detected from the lysates of the *Ec-EG* co-cultures and their respective single cultures were similar, and were significantly lower than the wild-type lysate (Figure 3G, H). This implies that thiamin is synthesized *de novo* in the *Ec-CE* and *Ec-CG* co-cultures. Further, LC-MS/ MS analysis of these samples confirmed the presence of thiamin in the lysates of the co-cultures (Figure S6). Taken together, these results show that the *Ec-CE* and the *Ec-CG* co-cultures grow due to *de novo* thiamin synthesis, whereas the *Ec-EG* co-cultures do not survive as they are unable to produce thiamin.

### *Vibrio anguillarum* thiamin mutants follow a similar pattern of exchange as *E. coli*

In order to determine whether the pattern of exchange of thiamin biosynthesis intermediates observed in *E. coli* is conserved across other microbes, we analyzed another gammaproteobacterium *Vibrio anguillarum* str. PF430-3 which is capable of *de novo* thiamin biosynthesis and salvage (Figure S1B). Similar to the previous experiment, *V. anguillarum ΔthiC, ΔthiE*, and *ΔthiG* mutant strains were grown individually and in pairwise co-cultures in thiamin-deficient M9 medium (Figure 4).

**Figure 4.**
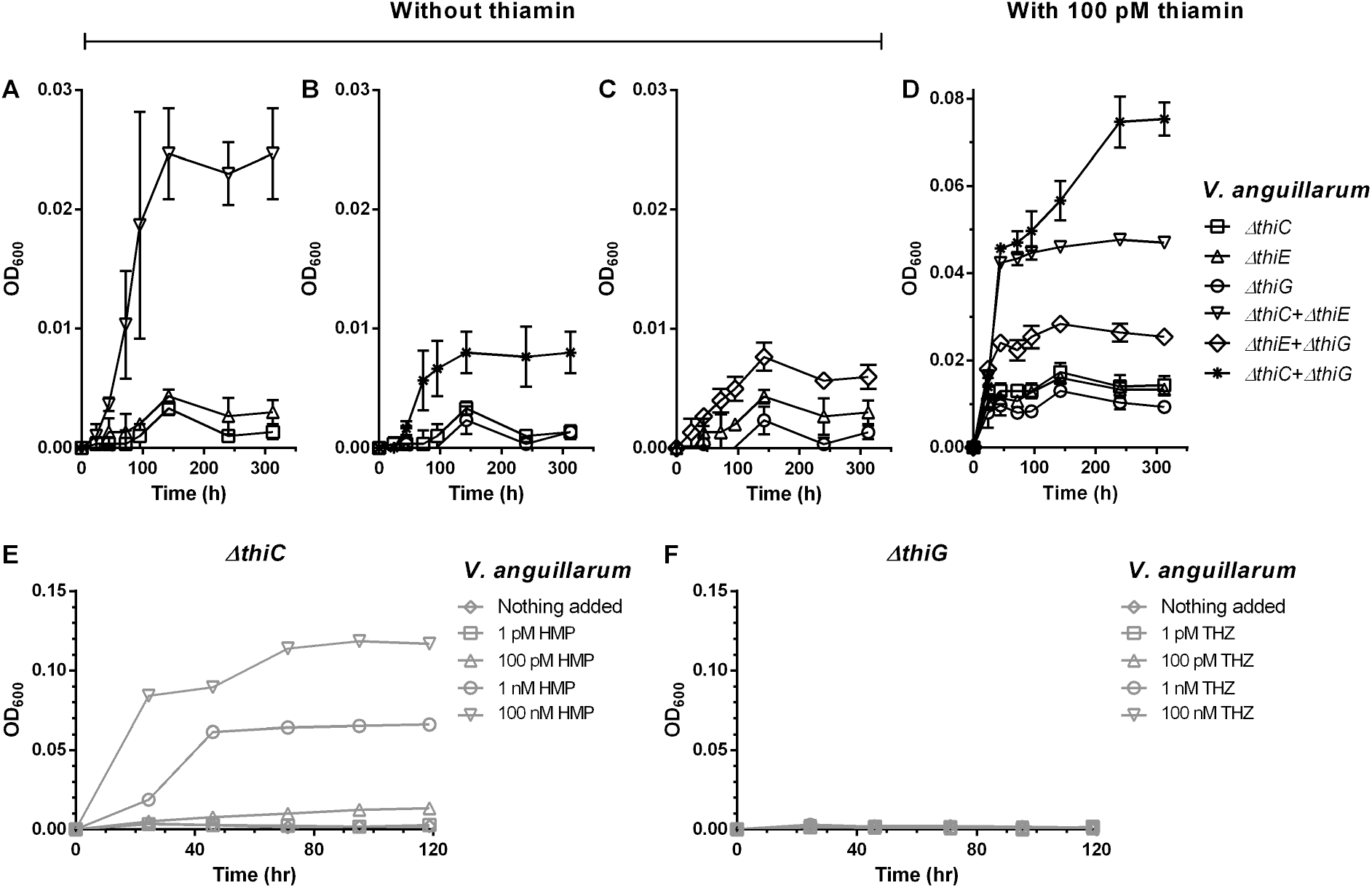
Thiamin biosynthesis mutants of *V. anguillarum* str. PF430-3 grow in pairwise co-cultures in thiamin-deficient M9 medium. Co-cultures of **(A)** the *VaΔthiC*-*VaΔthiE* strains, **(B)** the *VaΔthiC*-*VaΔthiG* strains, and **(C)** the *VaΔthiE*-*VaΔthiG* strains without thiamin supplementation. Co-cultures of **(D)** the *VaΔthiC, VaΔthiE*, and *VaΔthiG* strains supplemented with 100 pM thiamin. **(E)** HMP supplementation to the *VaΔthiC* mutant. **(F)** THZ supplementation to the *VaΔthiG* mutant. Symbols in panels **(D) and (F)** depict the following concentrations of HMP and THZ respectively: ◊ = nothing added, □ = 1 pM, Δ = 100 pM, ○ = 1 nM, ∇ = 100 nM. Means ± standard deviations from three independent experiments are plotted.

As *V. angiullarum* and *E. coli* are both gammaproteobacteria containing the same set of thiamin genes except the kinase *thiM*, we expect *V. anguillarum* to show a similar pattern of exchange (Figure S1). The *V. anguillarum ΔthiC, ΔthiE* and the *ΔthiG* single cultures showed no background growth in the P1 passage, and a concentration-dependent increase in growth starting with nanomolar concentrations of supplemented thiamin (Figure S7). For the co-cultures grown without supplemented thiamin, we observed that the *V. anguillarum CE* (*Va-CE*) co-culture showed significant growth followed by the *Va-CG* co-culture, while the *Va-EG* co-culture showed background growth, similar to what we observed in *E. coli* (Figures 4A-C). When supplemented with 100 pM thiamin, the *Va-CE, Va-CG* and *Va-EG* co-cultures grew significantly better than the single cultures, reiterating that the co-cultures were likely producing thiamin (Figure 4D). Interestingly, even though the *V. anguillarum ΔthiC* strain shows concentration-dependent increase in growth when supplemented with HMP similar to its *E. coli* counterpart, the *VaΔthiG* strain does not grow with exogenously added THZ (Figures 4E, F). This indicates that unlike the *E. coli ΔthiG* mutant, whose growth can be complemented by thiamin and THZ, the *V. anguillarum ΔthiG* mutant can be complemented only by thiamin. This may be attributed to the absence of the thiazole kinase gene *thiM* in *V. anguillarum*, which is considered to be a salvage enzyme that phosphorylates THZ to produce THZ-P for incorporation in thiamin biosynthesis in *E*.*coli* and other organisms (Figure 1, Figure S1).

### *V. anguillarum* and *E. coli* thiamin mutants exchange thiamin and its biosynthesis intermediates among themselves

Finally, to test whether the pattern of exchange that we observe occurs across different species, we created mixed co-cultures of the *E. coli ΔthiG* with *V. anguillarum ΔthiC* and *ΔthiE* strains in a pairwise manner. The *V. anguillarum ΔthiC*-*E. coli ΔthiG* (*VaC*-*EcG*) co-culture survived without thiamin supplementation as compared to the individual strains (Figure 5A). Surprisingly, the *VaE*-*EcG* co-culture also grew significantly better than the individual strains without thiamin supplementation (Figure 5B), even though the overall growth was lower than the *VaC-EcG* co-culture. This result is contrary to what was observed in the *Va-EG* or the *Ec-EG* co-cultures which did not grow beyond the background level. This indicates that THZ synthesized by the *V. anguillarum ΔthiE* strain might be available in the medium at a concentration that allows the *E. coli ΔthiG* to grow and produce thiamin and share it in return with *V. anguillarum ΔthiE*. The *VaC-EcG* and *VaE-EcG* co-cultures grow to similar OD_600_ with 100 pM of supplemented thiamin as expected (Figure 5C).

**Figure 5.**
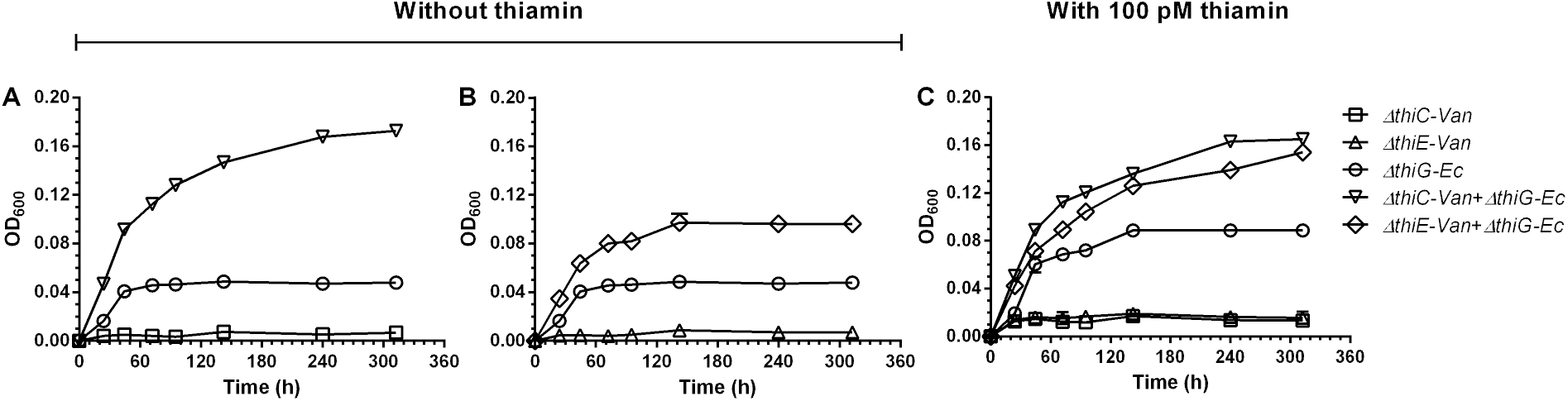
Mixed-species co-cultures of the thiamin biosynthesis mutants of *V. anguillarum* PF430-3 and *E. coli* K-12 MG1655 in thiamin-deficient M9 medium. Mixed co-culture of **(A)** the *VaΔthiC*-*EcΔthiG* strains and **(B)** the *VaΔthiE*-*EcΔthiG* strains without thiamin supplementation. **(C)** Co-culture of the *VaΔthiC* or *VaΔthiE* and *EcΔthiG* strains with 100 pM thiamin. Means ± standard deviations from three independent experiments are plotted.

To summarize, the *CE* and *CG* co-cultures of only *E. coli* or *V. anguillarum* strains can survive in the absence of externally supplemented thiamin, whereas the *EG* co-cultures cannot. These experiments implicate that in a community of thiamin auxotrophs, HMP and thiamin are shared more readily, but not THZ. Further, results obtained from the mixed co-cultures of *E. coli* and *V. anguillarum* strains suggest that even though THZ is picked up when present at higher concentrations, it might not be readily shared among microorganisms as the concentrations produced are too low to be salvaged.

### The ratio of the individual strains within the co-culture is determined by the exchange of thiamin and its biosynthesis intermediates

Our observations of the co-culture experiments thus far are based on the total OD_600_ of the co-cultures. To understand and quantify the contribution of each individual strain, we created the *thi*^*-*^ mutants fluorescently labeled with green fluorescent protein (GFP) and set up the pairwise *Ec-CE* and *Ec-CG* co-cultures in thiamin-deficient minimal media, where one of the strains in each co-culture was fluorescently labeled (Figure S2B, D). This approach allows us to quantify the amount of each strain in the co-culture using the two parameters – (i) the total OD_600,_ and (ii) the fluorescence of the co-culture which indicates the growth of the GFP-tagged strain. Briefly, we generated a standard curve of fluorescence versus OD_600_ for each individual strain, following which we set up the following pairs of cultures (GFP strain indicated with an asterisk) - *Ec-C*E* and *Ec-CE\** with the controls *Ec-C*E\** and *Ec-CE*, and a similar set for the *Ec-CG* co-cultures. We then noted the increase in the OD_600_ and fluorescence values over time and mapped the fluorescence signal of the co-culture to the standard curve of the corresponding GFP-tagged strain, allowing us to quantify its OD_600_ in the co-culture (Figure S8). The remaining untagged strain numbers were then calculated by subtracting this number from the total OD_600_, eventually yielding the ratios of the two strains over the course of the co-culture growth.

Our experiments and subsequent calculations showed that the quantities of the strains in the co-cultures change over a period of 24 h when no thiamin is exogenously provided (Figure 6 and Figure S9). The OD_600_ of the *Ec-C*E* co-culture increases over time as expected (Figure 6A). The fluorescence of the co-culture also increased, indicating that the quantity of the GFP-marked *EcΔthiC\** strain increased over time (Figure 6B). Next, we observed that for the *Ec-CE\** co-culture in the absence of thiamin, the fluorescence did not increase even though the OD_600_ value increased over time, reiterating that the *EcΔthiC* strain increased in numbers in the co-culture (Figures 6C and 6D).

**Figure 6.**
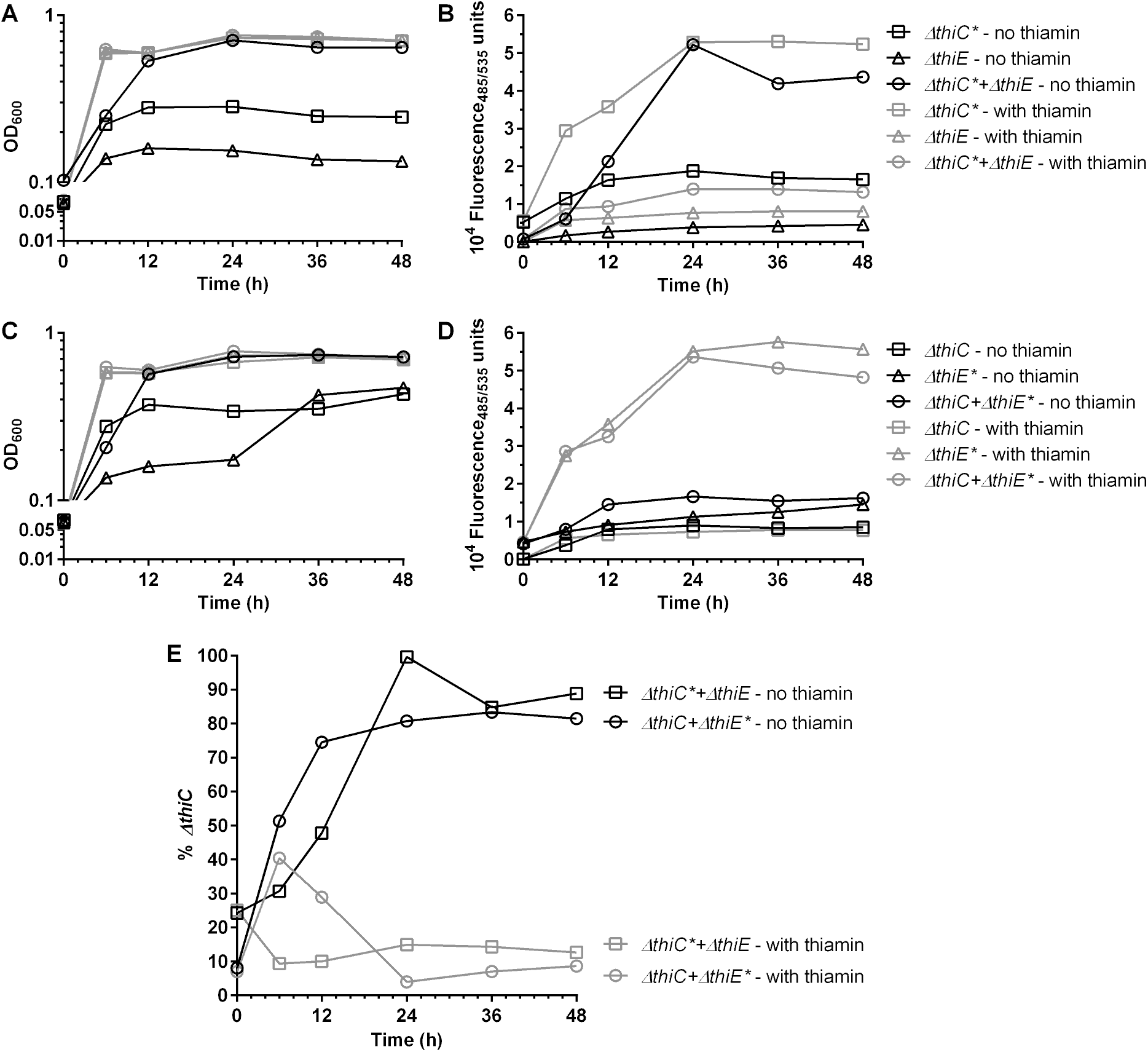
Growth phenotypes and fluorescence of the monocultures and the co-cultures of the thiamin mutant strains. The strains containing the *GFPmut2* cassette are marked with an asterisk. Black symbols = without thiamin, grey symbols = with thiamin. **(A)** OD_600_ and **(B)** fluorescence of *EcΔthiC\**-*EcΔthiE* co-cultures. **(C)** OD_600_ and **(D)** fluorescence of *EcΔthiC*-*EcΔthiE\** co-cultures. **(E)** Percentage of *EcΔthiC* cells in the *EcΔthiC\**-*EcΔthiE* co-cultures and the *EcΔthiC*-*EcΔthiE\** co-cultures. Average values from two independent experiments are plotted.

Interestingly, in the presence of thiamin, even though the OD_600_ of the *Ec-C*E* co-culture increased over time, and the fluorescence increase in the *EcΔthiC\** single culture cells was proportional to its growth as expected, the fluorescence of the *Ec-C*E* co-culture stayed constant over time (Figures 6A, B). This indicates that the ratio of the individual strains remains constant over time with respect to the starting ratio. Also, both the OD_600_ and the fluorescence of the *Ec-CE\** co-culture and the *EcΔthiE\** single culture increased over time, confirming that the numbers the two participating strains do not deviate in the co-culture in the presence of thiamin (Figures 6C, D).

Upon quantifying the *Ec-C*E* and *Ec-CE\** co-culture results, we found that in the absence of thiamin, the percentage of the *EcΔthiC* cells in the co-cultures increased over time, to attain an average ratio of ∼8:2 of *EcΔthiC*: *EcΔthiE* cells at 24 h (Figure 6E). Also, the presence of GFP does not alter the final ratios of the strains in the co-cultures as illustrated by both the *Ec-C*E* and *Ec-CE\** co-cultures showing similar ratios. Comparable ratios were obtained when the *Ec-C*G* co-cultures were similarly analyzed (Figure S9B). When the co-cultures were further transferred at the end of 24 h of growth to a fresh thiamin-deficient M9 medium in passage P3, the new ratios held constant over a period of 24 h (Figure S9A, C). Additionally, even after the continued growth of the P2 co-cultures for another ∼24 h, the ratios attained stayed constant (Figure 6E, S9B). We hypothesize that this change in the ratio of the two strains results from the exchange of HMP and thiamin which equilibrates after ∼24 h and subsequently stabilizes. However, in the presence of exogenously added thiamin, the exchange is no longer necessary and hence the ratios of the two strains remain mostly unaltered.

## Discussion

Thiamin, an essential nutrient for living organisms, assists enzymes in executing key decarboxylation reactions in primary metabolism. Several studies based on metagenomic analyses predict that thiamin and its building blocks HMP and THZ can be salvaged by both thiamin auxotrophs and prototrophs (16, 17, 41). In this study, we investigate the mechanism of thiamin synthesis and exchange within a microbial community through a molecular lens.

It has been reported that secondary transporters of thiamin such as PnuT which facilitate bidirectional transport of the vitamin are found more often in prototrophs, whereas the ABC family primary transporters such as ThiT which promote the uptake of thiamin are found more often in auxotrophs (16, 17). It has also been observed for both marine and gut microbial communities that some organisms in the community might be auxotrophic for the biosynthesis of both THZ and HMP, whereas certain others in the same community can produce both these intermediates, but lack the ability to combine them to form thiamin (16, 17). These observations reiterate that thiamin sharing is common among microorganisms.

To better understand the specifics of the exchange of thiamin and its intermediates in a community, we created synthetic co-cultures with bacterial strains with defined thiamin auxotrophy patterns. Our results from the *E. coli* and *V. anguillarum* co-cultures as well as mixed co-cultures of these two organisms show that thiamin and one of its biosynthesis intermediates HMP are commonly exchanged among microorganisms, whereas the exchange of the other intermediate THZ may occur less frequently and under specific conditions (Figure 7A). Our results show that the *Ec-EG* and *Va-EG* co-cultures do not grow, and we attribute this to the inability of THZ to be shared (illustrated in the schematic shown in Figure 7A and 1D). However, the mixed co-cultures i.e. in the *VaE-EcG* and the *VaC*-*EcG* co-cultures both show growth which indicates that there may be an exchange of THZ between these organisms (Figures 5A, B and 1D, E). Of these, the growth of the *VaE-EcG* co-culture was surprising and unexpected based on our previous results, and we reason out that there is only one possibility for how these two thiamin auxotroph strains may support one another’s growth – *V. anguillarum ΔthiE* supplies THZ to *E*.*coli ΔthiG*, which produces thiamin and in turn returns it to *V. anguillarum ΔthiE*, enabling it to grow and the co-culture to be sustained over 12 days (∼300 hrs) (Figure 5B). In the *VaC-EcG* co-culture, there are three possibilities as illustrated in schematic Figure 1E, briefly, (i) *VaΔthiC⟶* THZ *⟶ EcΔthiG, EcΔthiG ⟶* thiamin *⟶ VaΔthiC* (ii) *VaΔthiC⟶* THZ *⟶ EcΔthiG, EcΔthiG ⟶* HMP *⟶ VaΔthiC*, or (iii) *EcΔthiG ⟶* HMP *⟶ VaΔthiC, VaΔthiC ⟶* thiamin *⟶ EcΔthiG*. Based on the observation that the *VaE-EcG* is able to grow, it opens up the possibility for any of these to occur. However, as the OD_600_ of the *VaC-EcG* co-culture is significantly higher than the *VaE-EcG* co-culture, it is likely that the two co-cultures do not rely on the exchange of the same molecules (Figure 5A, B). Based on this observation, we hypothesize that the *VaC-EcG* co-culture may follow possibilities (ii) and (iii), and this needs to be investigated further.

**Figure 7.**
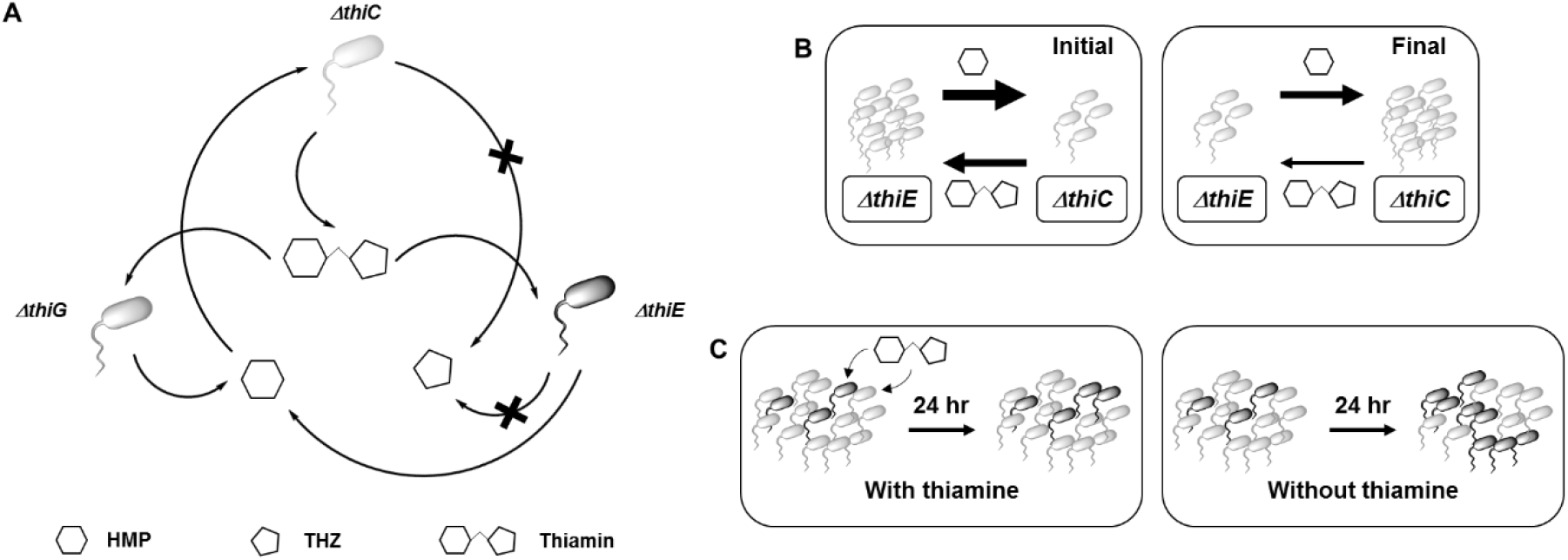
Proposed model for the exchange of thiamin biosynthesis intermediates in the co-cultures and effects of the exchange on the co-culture dynamics. **(A)** Probable molecular exchanges among the thiamin mutants. **(B)** Probable exchanges in the co-culture of the *ΔthiC*-*ΔthiE* strains without thiamin at the initial and the final stages (after 24 h) of the co-culture. Thickness of the arrows is proportional to the amounts of the respective nutrients being released. **(C)** Ratios of the two strains in the co-culture differ based on the presence or absence thiamin, black cells = *ΔthiC* strain, grey cells = *ΔthiE* or *ΔthiG* strain. Hexagon = HMP, pentagon = THZ, linked hexagon and pentagon = thiamin.

The unexpected growth of the *VaE-EcG* mixed co-culture in thiamin deficient media might be explained based on some of the characteristics of the co-culture inhabitants. The first possibility is that the cells of *VaΔthiE* lyse owing to the longer incubation time of ∼300 h as opposed to *EcΔthiE* cells in the *EcE-EcG* co-cultures which are grown for only 24 h. This results in the release of THZ in the medium, sufficient amount of which then accumulates and is salvaged by the *EcΔthiG* cells, and thus the *VaE-EcG* co-culture survives. But had this been the case, the *Va-EG* co-culture which showed no growth for ∼300 h should have also survived (Figure 4C). We hypothesize that this inability to grow is because unlike *E. coli* which harbors ThiM, *V. anguillarum* lacks this enzyme that is essential for the conversion of THZ to THZ-P which is then routed into thiamin biosynthesis. Thus in respective co-cultures, adequate quantity of THZ is derived from *VaΔthiE* cells either through lysis or release. But as the *VaΔthiG* cells lack ThiM and thus cannot make THZ-P and subsequently thiamin, the *Va-EG* co-culture does not survive. On the other hand, as the *EcΔthiG* cells convert THZ to THZ-P and synthesize thiamin and provide it back to the *VaΔthiE* cells, the *VaE-EcG* co-culture survives in the absence of any exogenous thiamin.

When calculating the ratios of the two strains in the co-culture, we noted that the *EcΔthiC* strain increases in the co-culture over time and the ratios of *EcΔthiC*: *EcΔthiG* and *EcΔthiC*: *EcΔthiE* finally stabilize at ∼8:2. The role of the *EcΔthiG or EcΔthiE* strains in both co-cultures is to provide HMP, whereas that of *EcΔthiC* is to produce thiamin. Let us take the instance of the *Ec*-*CE* co-culture. The *EcΔthiE* strain (present in a higher amount at the start) produces HMP and supplies it to the *EcΔthiC* strain. The *EcΔthiC* strain produces thiamin, and is now self-sufficient. However, as it grows and replicates, it will require more thiamin and hence, it needs a small but continuous supply of HMP, and hence, it provides the *EcΔthiE* strain with just enough thiamin such that some *EcΔthiE* cells continue to survive in the co-culture (Figure 7B). Thus, we can conclude that the strain that produces thiamin, which both strains need, plays a more significant role in the co-culture.

We also observed that when thiamin is supplemented to the co-cultures of the *E. coli thi-* mutants, the ratios of the two strains in the co-cultures do not deviate much from the starting ratios (Figure 7C). This suggests that when a nutrient is available in plenty in a community of auxotrophs, they may not interact with each other. But when the nutrient is unavailable or scarce, a crosstalk that allows for the microorganisms to collectively build and share the nutrient may evolve, which will result in subsequently shaping the composition of and relative abundance of members of the community. Indeed, it has been shown that the seasonal blooms of marine microorganisms which either produce or utilize thiamin alter concentrations of thiamin biosynthesis intermediates in seawater and when the microbial numbers are low, the overall concentrations of the intermediates remain at an equilibrium (32). Such changes in the community composition have also been reported earlier for synthetic co-cultures based on their differential ability of nutrient exchange or uptake (4, 5, 39).

Finally, we hypothesize that the reason for HMP being exchanged more readily as compared to THZ among auxotrophs is that the biosynthesis of THZ is more intricate as compared to the biosynthesis of HMP. THZ is assembled by *thiG* (or thi4 in eukaryotes) using three distinct intermediates from different pathways, one of which includes a series of intricate sulfur-transfer reactions, whereas HMP biosynthesis by *thiC* is a rearrangement of a single intermediate. Thus, the biosynthesis of HMP may have a lower metabolic expense as compared to THZ, making it easier for organisms to share HMP rather than THZ. Interestingly, one study reports that the ratio of the *thiC:thiG+thi4* genes in marine microbes is in the 0.06-0.28 range, always less than one (41). Congruently, another study reported higher concentrations of HMP than thiamin in surface waters of the Sargasso Sea, and that the abundance of *thiC* genes was lesser than the *thiG* genes at depths of 0, 40 and 80 m (23). Even beyond marine ecosystems, there is a propensity for HMP exchange within the human gut microbiome (HGM) as well, wherein out of the 2,228 reference genomes studied, 199 were HMP auxotrophs, whereas only 114 were THZ auxotrophs (17). These studies, taken together with our observations, point to HMP and possibly other pyrimidine intermediates that can yield HMP via salvage as key nutrients in determining the dynamics of nutrient exchange and subsequently microbial abundance (22, 31, 32).

## Conclusion

In this study, we have designed a unique co-culture system based on the exchange of intermediates derived from the same metabolic pathway, in this case vitamin B_1_ biosynthesis. We conclude that the sharing of vitamin B_1_ and its intermediates is modulated by the availability of as well as the presence of biosynthesis and transporter proteins in cells. Exchange forms the basis of building an interacting community of microbes, but may also be a feasible mechanism to halt interactions or limit the success of portions of a community, e.g. provision of thiamin rather than HMP to prevent dominance of pyrimidine auxotrophs. Finally, our investigations at the molecular level underscore the specific role of metabolite exchange in determining, stabilizing and sustaining the collective metabolism and composition of our microbial co-cultures, making it possible to create defined communities for synthetic biology and biotechnological applications in the future.

## Materials and methods

### Chemicals and reagents

All the chemicals used were obtained either from TCI, HiMedia or Sigma unless otherwise specified. The enzymes used were obtained from TaKaRa.

### Strains and plasmids

The *E. coli* K-12 MG1655 strain containing pKD46 and the plasmids pKD3 and pProEX-Hta were a gift from Dr. Nishad Matange at IISER Pune. The plasmids pCA24N-*EcthiC*, pCA24N-*EcthiG*, and pCA24N-*EccobT* were obtained from the ASKA collection hosted at IISER, Pune. The *E. coli* KL-16 strain harboring the *GFPmut2-kan*^*R*^ cassette was a gift from Dr. Deepa Agashe at NCBS, Bangalore.

### Generating single gene knockouts in E. coli

All the single knockout mutants of *E. coli* K-12 MG1655 used in the study were generated using recombination by λ Red Recombineering system (36, 43). The primers sequences used for generating the gene knockouts and for their verification are listed in the supplementary table 1. For generating the strains marked with GFP, we flipped out the *kan*^*R*^ cassette from the *thi*^*-*^ mutants of *E. coli* K-12 MG1655. We then cloned and inserted the *GFPmut2::kan*^*R*^ cassette from the *E. coli* KL-16 strain into the *thi*^*-*^ mutants, after *aidB* gene, in the reverse orientation with respect to the *aidB* gene. This gave us the following *E. coli* mutants – *thiC aidB1633::GFPmut2-kan*^*R*^ (*EcΔthiC\**), *thiE aidB1633::GFPmut2-kan*^*R*^ (*EcΔthiE\**) and *thiG aidB1633::GFPmut2-kan*^*R*^ (*EcΔthiG\**). The *GFPmut2-kan*^*R*^ insertions were carried out using the same λ Red Recombineering system mentioned above.

### Cloning of EcthiE, and transformations of the pCA24N plasmids

The *thiE* gene from *E. coli* K-12 MG1655 was cloned in pProEx-Hta vector using restriction-free cloning method as previously described (44). The empty pProEx-Hta vector and the pProEx-Hta-*EcthiE* vector were then chemically transformed into the *E. coli* K-12 MG1655 *ΔthiE::kan*^*R*^ strain for complementation analysis. The pCA24N-*EccobT* was chemically transformed into *E. coli* K-12 MG1655 *ΔthiC::kan*^*R*^, *ΔthiE::kan*^*R*^, and *ΔthiG::kan*^*R*^ strains for complementation analysis. The pCA24N-*EcthiC* and the pCA24N-*EcthiG* plasmids were chemically transformed into the *E. coli* K-12 MG1655 *ΔthiC::kan*^*R*^ and *ΔthiG::kan*^*R*^ strains respectively.

### Growth conditions and media

*E. coli* K-12 MG1655 cells were grown either in Luria-Bertani Miller (LB) or in M9 salts minimal medium at 37°C, 180 rpm (45). Whenever necessary, the medium was supplemented with various components in small defined amounts as mentioned.

### Primary culture set-up (LB and P1 cultures) for E. coli

*E. coli* K-12 MG1655 WT, *ΔthiC, ΔthiE*, and *ΔthiG* mutants were grown in LB aerobically at 37°C, 180 rpm, for 6-8 hours. The cultures were centrifuged at 6500 rpm for 1 minute and the pellets were washed thrice with 1X M9 salts by re-suspending them using a vortex for each wash. This step was used to make sure that the cells do not carry-over any residual nutrients from LB. These cells were used to start P1 cultures (first subcultures in minimal medium) in [M9 + NH_4_Cl + Glucose + Inosine (50 µM)] medium, in 4 mL medium in 25 mL test-tubes, at a starting OD_600_ of 0.05, and were incubated aerobically at 37°C, 180 rpm, for 16-18 hours. Cells grown in P1 were centrifuged at 6500 rpm for 1 minute and the pellets were washed thrice with 1X M9 salts.

For the *thi*^*-*^ mutant rescue experiments, the mutants with or without pProEx-Hta or pCA24N plasmids harboring the genes mentioned were grown similarly in P1, supplemented with or without thiamin (20 µM).

### Pairwise co-culture set-up of E. coli mutants in P2

*E. coli* cells washed after P1 were used to start their co-cultures in P2 (second subcultures in minimal salts medium – composition described above) at a starting OD_600_ of 0.1, in a 96-well plate with lid, with 200 µL medium in each well, and were incubated aerobically at 37°C, ∼240 rpm orbital shaking, for 24-96 hours, with OD_600_ reading and fluorescence reading at excitation/ emission values of 485/ 535 after every shaking cycle of ∼300 sec, with upper lid at a temperature 2°C higher than 37°C (in EnSight) to avoid condensation, inside a plate reader – either Tecan or EnSight respectively. Alternately, the cells were grown in 25 mL test tubes with 4 mL medium each at 37°C, 180 rpm, in a shaker incubator. The media used were supplemented with various nutrients as and when required in the concentrations mentioned, for both the thiamin requirement and the HMP and THZ feeding studies. The two different mutants used for co-cultures were inoculated either singly as a control or in their co-culture at the ratios of 1:9 or 1:1 or 9:1. Both the single cultures as well as the co-cultures were inoculated at a starting OD_600_ of 0.1, that is, say, for the co-cultures with the 1:9 ratio, the two mutants were mixed at a starting OD_600_ of 0.01 and 0.09 of the individual mutants, respectively. When using the alternate carbon sources, glucose (final concentration in the medium - 22.2 mM) was replaced with 33.3 mM Na-succinate or 44.4 mM Na-pyruvate, to keep the amount of carbon fed the same for all media.

### Re-inoculation experiments in P3

For the re-inoculation experiments, the cells from the single cultures or the co-cultures were harvested at the end of 24 h, washed thrice with 1X M9 salts, re-inoculated in the fresh M9 minimal medium devoid of thiamin, at a starting O.D._600_ of 0.1, and incubated further at 37°C, 180 rpm, in 25 mL test tubes with 4 mL medium each.

### OD_600_ and fluorescence correlation

For the OD_600_ and fluorescence correlation, the cells were grown in all four possible pairwise combinations of the *EcΔthiC::kan*^*R*^ strain or the *EcΔthiC\** strain with the *EcΔthiE::kan*^*R*^ strain or the *EcΔthiE\** strain. The values of the total fluorescence of the co-cultures carrying a single fluorescent strain were then normalized using the co-culture of the *EcΔthiC::kan*^*R*^ strain with the *EcΔthiE::kan*^*R*^ strain as a control, and the OD_600_ value of that fluorescent strain was calculated using the fluorescence v/s OD_600_ correlation of that strain. From this exercise, we obtained the ratios of the strains in the co-cultures of the *EcΔthiC::kan*^*R*^ strain with the *EcΔthiE::kan*^*R*^ strain and those of the *EcΔthiC::kan*^*R*^ strain with the *EcΔthiG::kan*^*R*^ strain at different time points.

### Thiochrome assay to detect the presence of thiamin from lysates

2 mL cells each from the single cultures and co-cultures were harvested at different time points and lysed in 125 µL of 1X PBS. The lysates were centrifuged at 14000 rpm, 4°C, and 100 µL each of the clarified lysate or spent medium were used to detect the presence of thiamin, TMP and TDP using HPLC-FLD and LC-MS/MS. The thiochrome assay was carried out as per a protocol previously described (42). Thiochrome formed was detected using HPLC-FLD (Agilent) on a C-18 reverse phase column (Phenomenex – Gemini), at 25°C. The solvents used were MilliQ in line A, methanol in line B, and 10 mM CH_3_COONa, pH 6.6 in line C. 100 µL of the standards or the samples were injected in HPLC for analysis. A standard curve on HPLC was generated using various concentrations of thiamin.HCl, TMP and TDP standards. The flow rate was maintained at 0.5 mL/ min. The HPLC and LC-MS/MS method used was 0 min: 100% A; 4 min: 90% A, 10% B; 20 min: 15% A, 25% B, 60% C; 24 min: 15% A, 25% B, 60% C; 30 min: 100% A; 44 min: 100% A. All samples were ran in positive ion mode with ESI method of ionization on SciEX-X500LR system for LC-MS/MS analysis.

### *V. anguillarum* cultures and experiments

*Vibrio anguillarum* PF430-3 (46, 47) wild-type and mutants *ΔthiC, ΔthiE*, and *ΔthiG* (41) were used in experiments. All were re-isolated from cryopreserved stock using marine broth agar plates and liquid medium (48) with estuarine surface water from MODMON Neuse River Estuary monitoring station 180 (49) as the base medium. Cells from liquid ZoBell cultures in late exponential or early stationary phase were washed and centrifuged (9,000 g, 3 min) thrice with 1X M9 medium without thiamin and then resuspended in M9 medium without thiamin. Absorbance at 600 nm (opitical density, OD) was measured using a spectrophometer (GENESYS 30, Thermo). Based the OD of resuspended cell cultures, 0.001 OD of washed cells was added (final density) at the start of each experiment.

Co-cultures of PF430-4 strains were started by adding 0.001 OD (final conc.) of each strain to M9 medium. Cultures were grown in clear sterile polystyrene tubes, and incubated in the dark at 20°C with daily homogenization by repetitive inversion. Thiamin hydrochloride, HMP, THZ used in experiments were purchased from TCI, Alfa Aesar, Fisher Scientific at ≥98% HPLC purity. Fresh solutions of vitamins were prepared under reduced light in a laminar flow hood autoclaved MilliQ water as the dilutant. Solutions were kept on ice while setting up experiments.

*E. coli JW5549 ΔthiG761::kan* (Keio Collection) was re-isolated as for PF430-4 but using M9 as the base for ZoBell solid and liquid media. Cells were washed and resuspended in M9 medium without B1 as for PF430-4. Co-cultures of PF430-4 *ΔthiC* or *ΔthiE* and *E. coli JW5549 ΔthiG761::kan* were initiated by adding 0.001 OD of each cell type to M9 medium without amended B1. Growth conditions were the same as for PF430-4.

## Acknowledgements

We acknowledge Deepa Agashe and Nishad Matange for providing specific *E. coli* strains and plasmids. We thank Gayathri Pananghat, Manjula Reddy, and Nishad Matange for insightful discussions. We thank PerkinElmer-IISER Pune Centre of Excellence, IISER Biology LC-MS facility, and Sandanaraj Britto and Thomas Pucadyil labs for instrumentation. We also thank Yamini Mathur, Yashwant Kumar, and Ateek Shah for their critical inputs. We acknowledge Saswata Nayak, Rituparna Ghosh, Mir Nasir Ahmed, and Gina Welsing for their contribution.

