## Supplementary information for "The exchange of vitamin B_1_ and its biosynthesis intermediates in synthetic microbial communities shapes the community composition and reveals complexities of nutrient sharing"

### Figures – Supplemental material

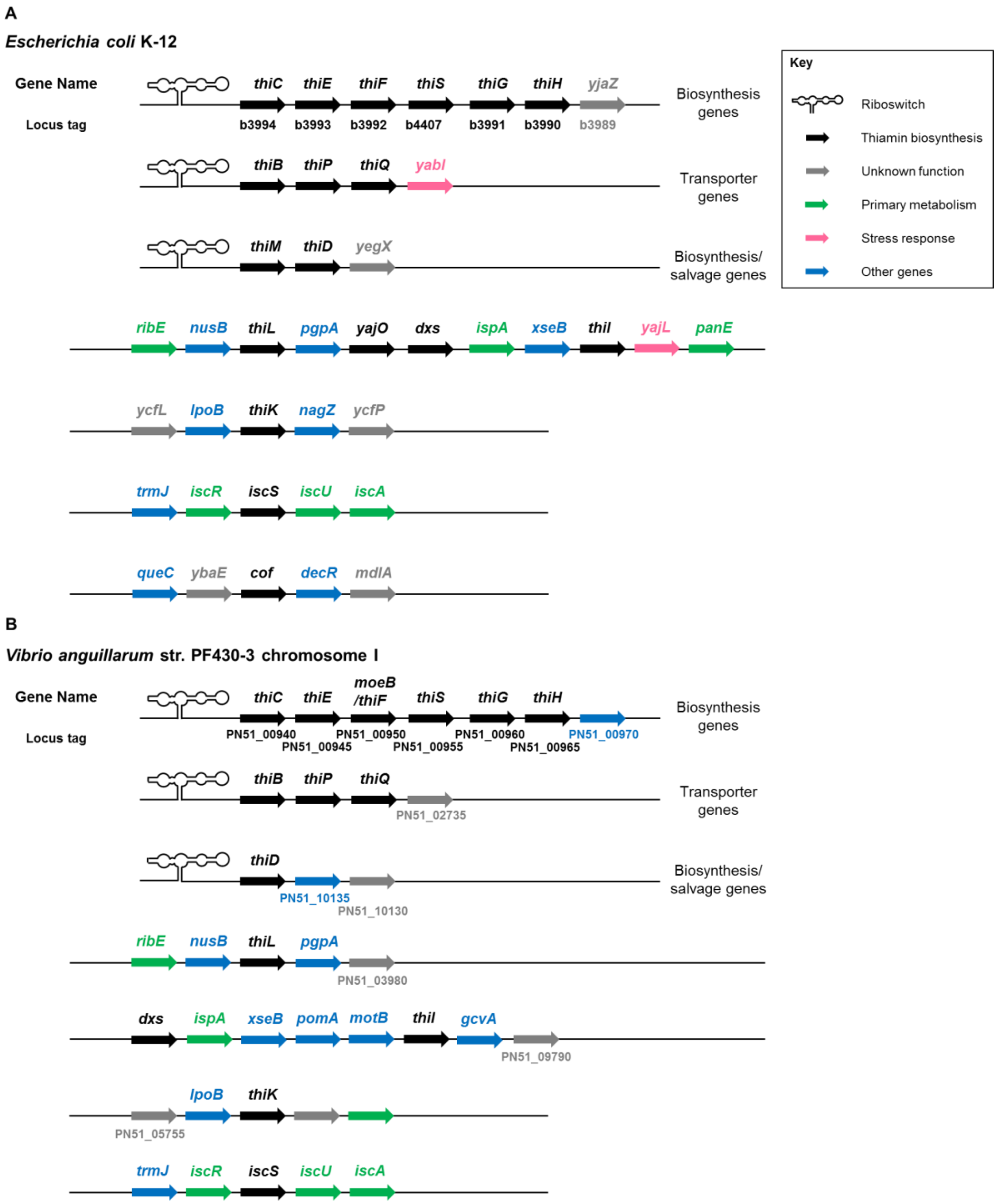


**Figure S1.** Arrangement of the thiamin biosynthesis genes on the genome of **(A)** *E. coli* K-12 MG1655 and **(B)** *V. anguillarum.* Note that other than the kinase gene *thiM* that is present in *E. coli* and is missing in *V. anguillarum*, the two organisms contain all other thiamin biosynthesis genes.

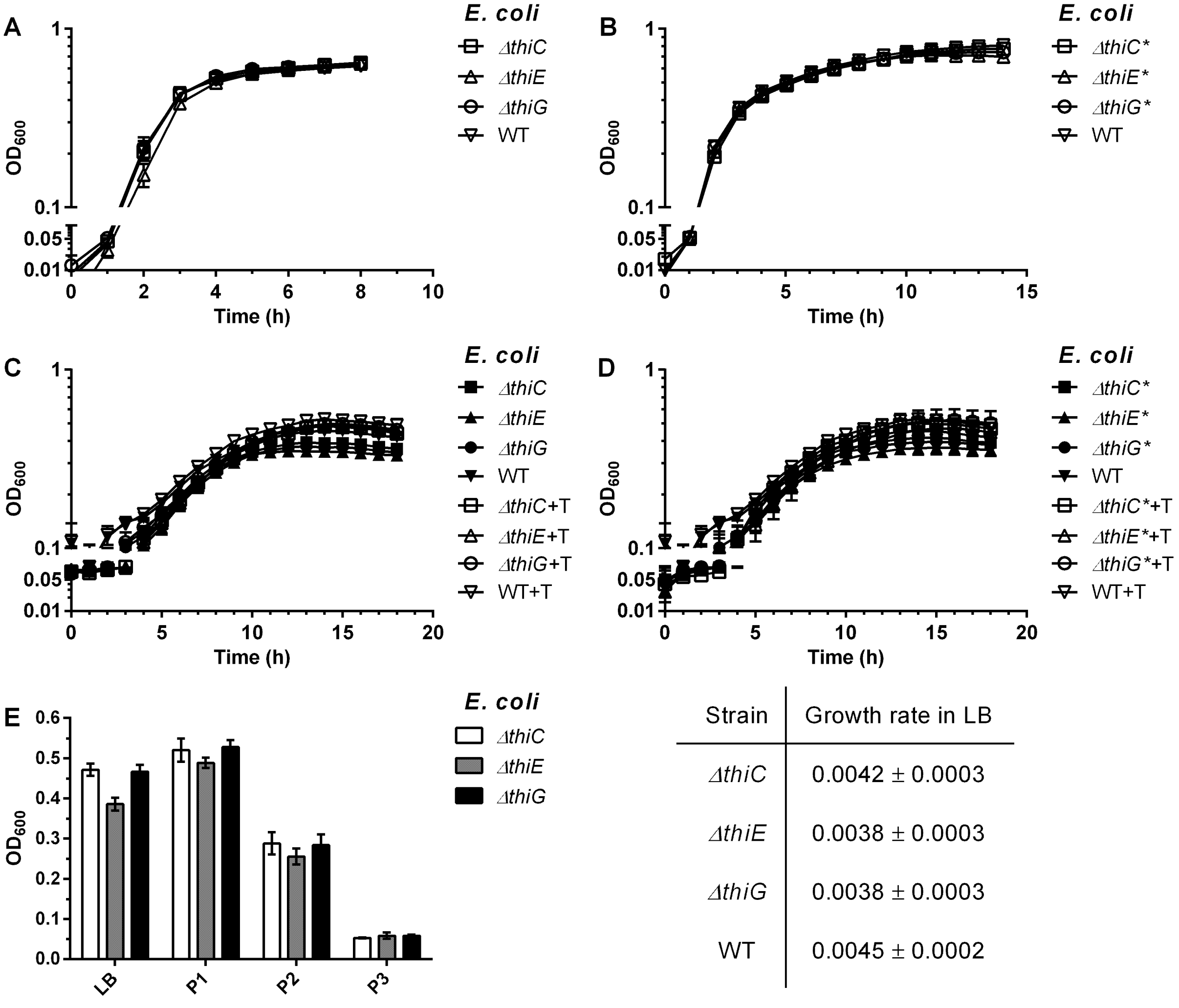


| Strain | Growth rate in LB |
| --- | --- |
| *ΔthiC* | 0.0042 ± 0.0003 |
| *ΔthiE* | 0.0038 ± 0.0003 |
| *ΔthiG* | 0.0038 ± 0.0003 |
| WT | 0.0045 ± 0.0002 |

**Figure S2.** Growth phenotypes of *E. coli* thiamin biosynthesis mutants in LB and M9 medium. Means ± standard errors of the means from three independent experiments are plotted. **(A)** Growth phenotypes of non-GFP tagged mutants in LB. **(B)** Growth phenotypes of GFP-tagged mutants in LB. **(C)** Growth phenotypes of non-GFP tagged mutants in P1. **(D)** Growth phenotypes of GFP tagged mutants in P1. **(E)** Growth phenotypes of non-GFP tagged mutants in P1, P2, and P3 at the end of 18h, 24 h, and 24 h, respectively, and started at $OD_{600}$ of 0.05, 0.1, and 0.1 respectively.

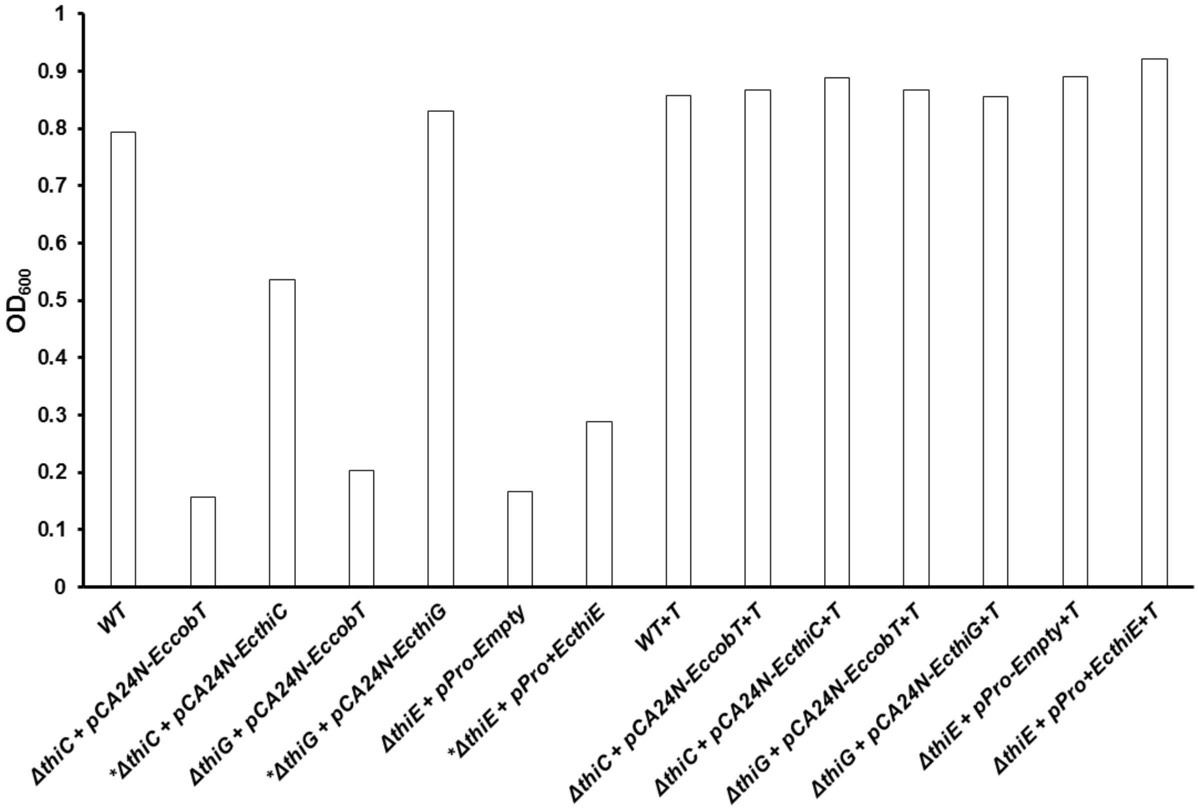


**Figure S3.** Growth of *E. coli* thiamin biosynthesis mutants is restored in P1 after 18 h of growth when complemented with a plasmid carrying the missing genes (indicated by *). The pCA24N plasmid containing the *EccobT* gene (from a $B_1$-unrelated pathway) was used as a negative control plasmid for the *Ec⊿thiC* and the *Ec⊿thiG* mutant rescue experiments. The pPro empty plasmid was used as a negative control for the *Ec⊿thiE* mutant rescue experiment. For the wild-type and thiamin-supplemented controls, results of a single experiment are plotted; whereas for the rest, average values from two independent experiments are plotted.

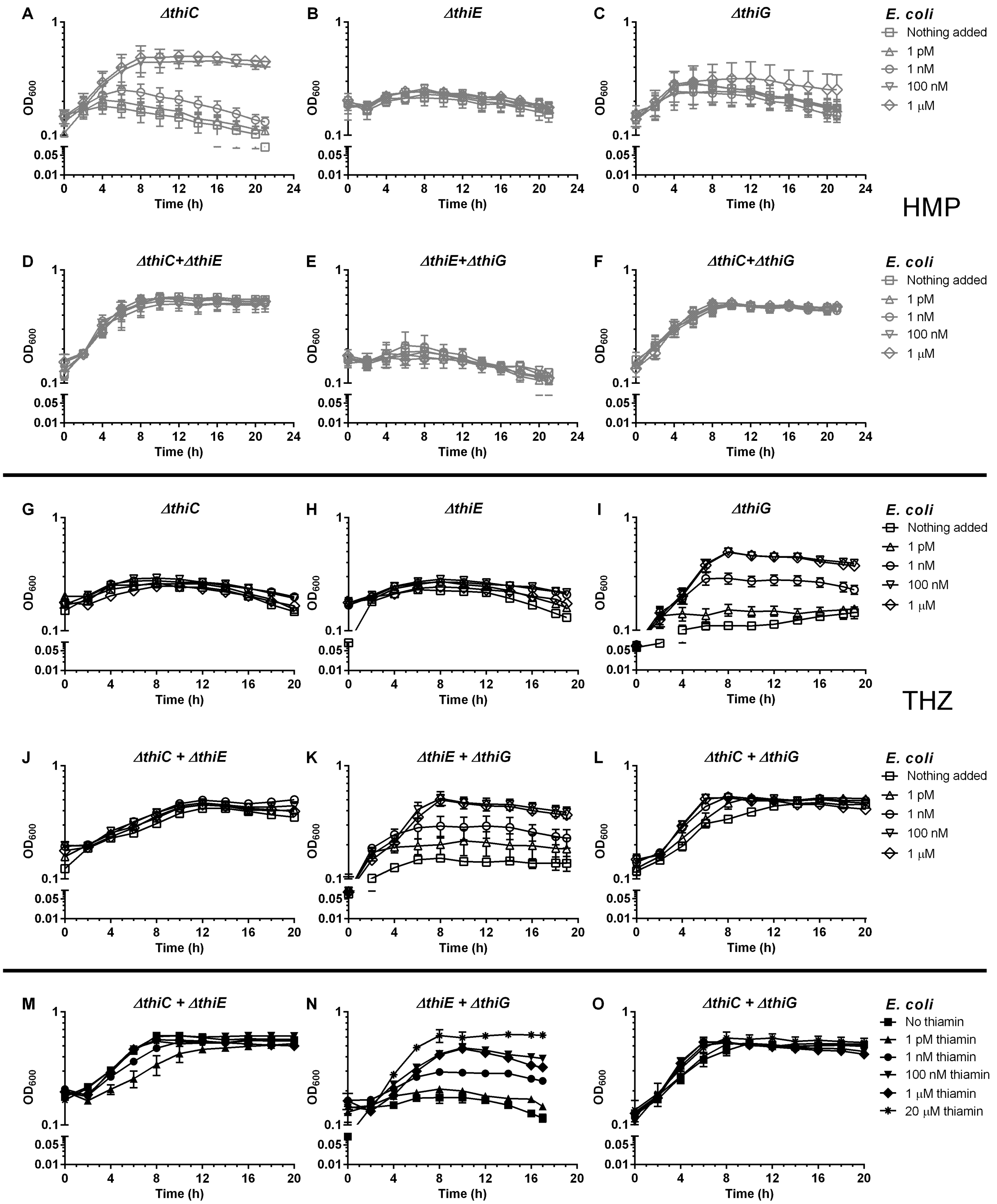


**Figure S4.** Supplementation of HMP, THZ and thiamin to the monocultures and the co-cultures of *E. coli* thiamin mutants in P2. **(A-F)** HMP supplementation, **(G-L)** THZ supplementation, **(M-O)** thiamin supplementation. Empty grey symbols – HMP supplementation, empty black symbols – THZ supplementation, and filled black symbols – thiamin supplementation. Means ± standard errors of the means from three independent experiments are plotted. HMP supplementation to **(A)** the *ΔthiC* strain, **(B)** the *ΔthiE* strain, **(C)** the *ΔthiG* strain, **(D)** the *Ec-CE* co-culture, **(E)** the *Ec-EG* co-culture, and **(F)** the *Ec-CG* co-culture. THZ supplementation to **(G)** the *EcΔthiC* strain, **(H)** the *EcΔthiE* strain, **(I)** the *EcΔthiG* strain, **(J)** the *Ec-CE* co-culture, **(K)** the *Ec-EG* co-culture, and **(L)** the *Ec-CG* co-culture. Thiamin supplementation to **(M)** the *Ec-CE* co-culture, **(N)** the *Ec-EG* co-culture, and **(O)** the *Ec-CG* co-culture.

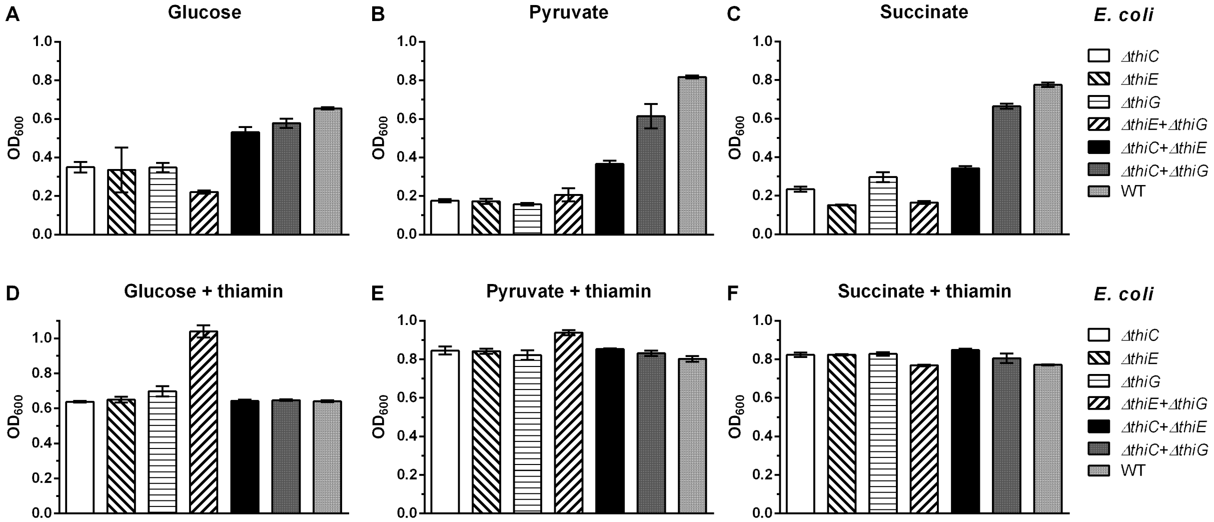


**Figure S5.** Growth phenotypes of the monocultures and the co-cultures of *E. coli* thiamin mutants under different physiological conditions.

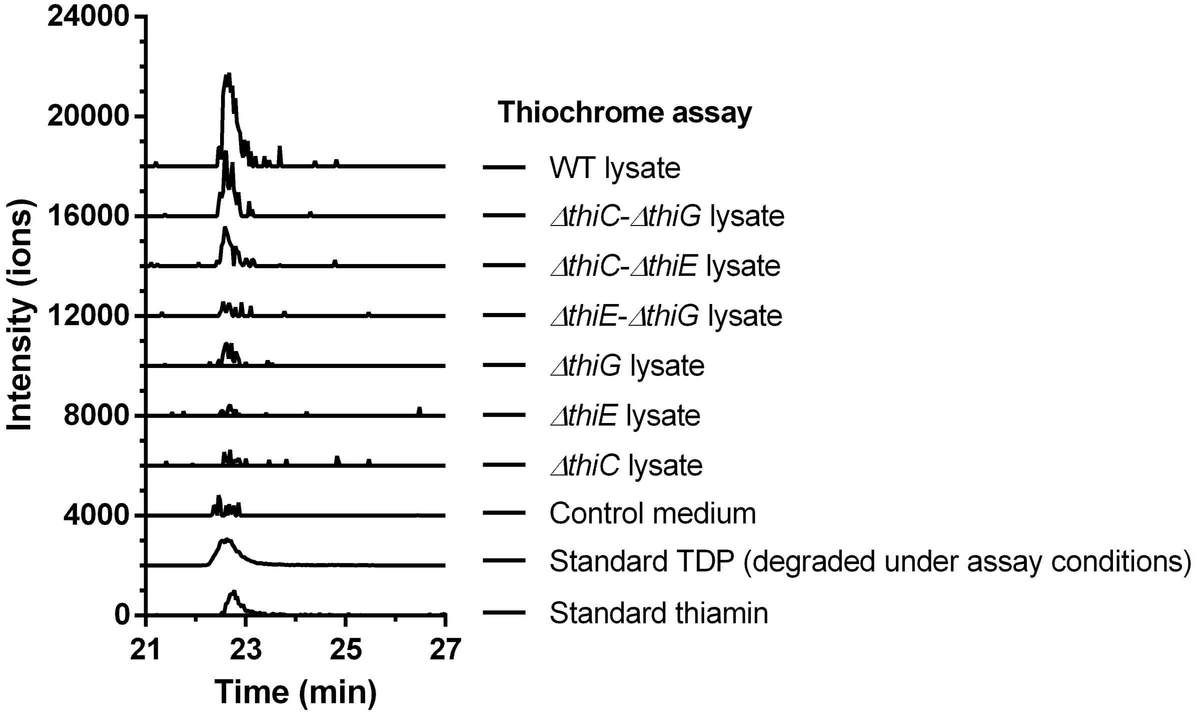


**Figure S6.** LC-MS analysis of the lysates of the co-cultures and the monocultures of *E. coli* for the detection of thiamin using the extracted mass of thiamin thiochrome (m/z 263.0888). The peak observed at 22.75 min for thiochrome in the *Ec-CE* and the *Ec-CG* co-culture is higher than those observed in the single cultures and the *Ec-EG* co-culture. The data is normalized to volumes of the lysates and standards injected.

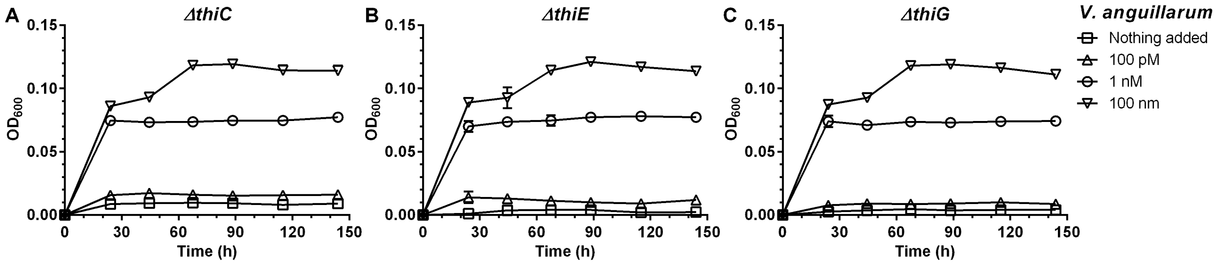


**Figure S7.** Supplementation of thiamin to the monocultures of *V. anguillarum* thiamin mutants in P1. Thiamin supplementation to **(A)** the *VaΔthiC* strain, **(B)** the *VaΔthiE* strain, and **(C)** the *VaΔthiG* strain. Means ± standard deviations from three independent experiments are plotted.

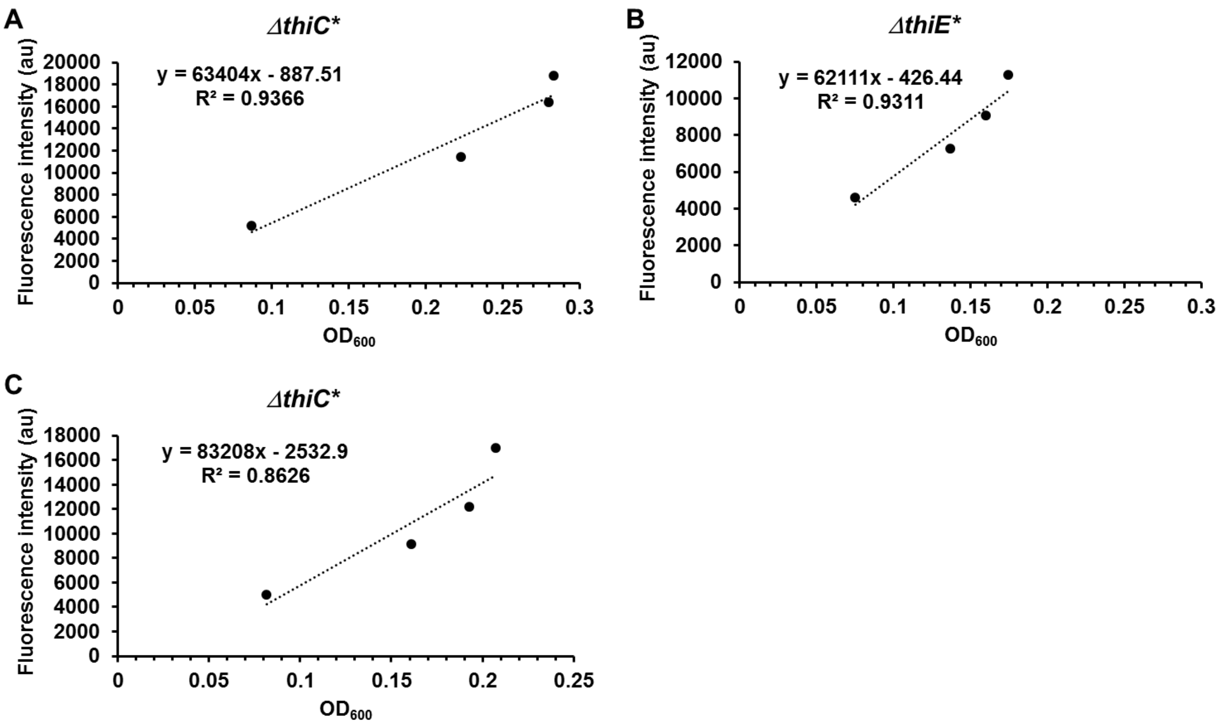


**Figure S8.** Fluorescence vs $OD_{600}$ correlation for the thiamin biosynthesis mutants of *E. coli* grown in P2 for data in figures 6 **(A and B)** and S9 **(A-C)**. Correlation for **(A)** the $\Delta$*thiC* strain in the *Ec-C*E* co-culture, **(B)** the $\Delta$*thiE* strain in the *Ec-CE** co-culture, and **(C)** the $\Delta$*thiC* strain in the *Ec-C*G* co-culture. Average values from two independent experiments have been plotted.

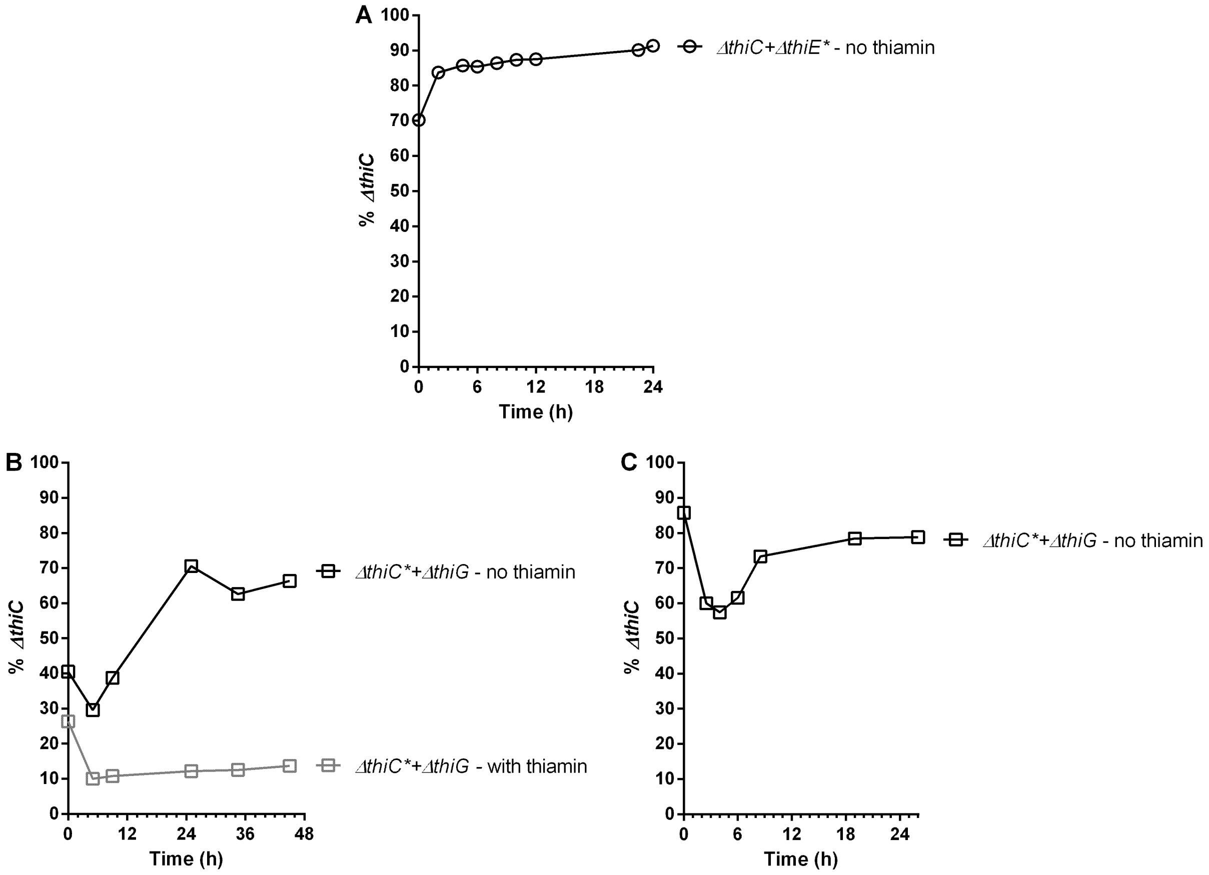


**Figure S9.** Percentage of *E. coli ΔthiC* strain in **(A)** the *Ec-CE** co-culture re-inoculated in fresh thiamin-deficient M9 medium in the third passage (P3) after 24 h of growth in P2 (data shown in figure 6E), **(B)** the *Ec-C*G* co-culture grown in P2, and **(C)** the *Ec-C*G* co-culture re-inoculated in fresh thiamin-deficient M9 medium in the third passage (P3) after 24 h of growth in P2. Average values from two independent experiments have been plotted.

#### Tables – Supplemental material

| Sr. No. | Primer name | Purpose | Primer Sequence (5' → 3') |
|---|---|---|---|
| 1. | *thiC::kan*$^R$_for | Forward primer to replace *E. coli thiC* with kanamycin cassette | TCATCCGTCGTCTGACAAGCCACGTCCTTAACTTTTTGGAATGAGCTATGGTGTAGGCTGGAGCTGCTTC |
| 2. | *thiC::kan*$^R$_rev | Reverse primer to replace *E. coli thiC* with kanamycin cassette | AGGTACAGGAGGAAAATCAGGCTGATACATCACGCTTCCTCCTTACGCAGCATATGAATATCCTCCTTAG |
| 3. | *thiE::kan*$^R$_for | Forward primer to replace *E. coli thiE* with kanamycin cassette | TTCCGTGCCAGAGGCGGAGAAATCTACCTGCGTAAGGAGGAAGCGTGATGGTGTAGGCTGGAGCTGCTTC |
| 4. | *thiE::kan*$^R$_rev | Reverse primer to replace *E. coli thiE* with kanamycin cassette | CCCGTCCAGAGCGATATCGTCGAGCAGGATCATTCATCGCCAACTCCTGCCATATGAATATCCTCCTTAG |
| 5. | *thiG::kan*$^R$_for | Forward primer to replace *E. coli thiG* with kanamycin cassette | GGATGGCGACCAGATCCTGCTTTTTCAGGTTATTGCAGGGGGTTGAAATGGTGTAGGCTGGAGCTGCTTC |

| | | | |
|---|---|---|---|
| 6. | *thiG::kan$^R$*_rev | Reverse primer to replace *E. coli thiG* with kanamycin cassette | CAGTTGTCGCCAGCGATCGCTGAAGGTTTTCATGCCGATGCCTCCAGAAACATATGAATATCCTCCTTAG |
| 7. | *thiC::kan$^R$*_conf_for | Forward primer to confirm *ΔthiC* mutation | CGAAGGGAACAAGAGTTAATCTGC |
| 8. | *thiE::kan$^R$*_conf_for | Forward primer to confirm *ΔthiE* mutation | ATTTTTGCTCCATGTGTGGG |
| 9. | *thiG::kan$^R$*_conf_for | Forward primer to confirm *ΔthiG* mutation | CAAACTGTTCACGAACTACTGGAG |
| 10. | *kan$^R$*_conf_rev | Reverse primer to confirm *thi$^-$* mutants | TAATCAGCACCTGGCTGTCG |
| 11. | *aidB_GFPmut2*_FP | Forward primer to insert *GFPmut2* cassette | GAATGATTTATTGCTGCGGGCGACGGGGGGAGTGTGTGTGTAAGCGTATACCAACTCAGCTTCC |
| 12. | *aidB_GFPmut2*_RP | Reverse primer to insert *GFPmut2* cassette | CAATTTTCACATATTTCATTTAGTTAATCGAAACCAGCGTCGCATCAGTCGATGAGCTCGAGATAT |

| | | | |
|---|---|---|---|
| 13. | *Ec_thiE*_PCR1_FP | Forward primer to clone *E. coli thiE* gene from *E. coli* genome | ATGTATCAGCCTGATTTTCC |
| 14. | *Ec_thiE*_PCR1_RP | Reverse primer to clone *E. coli thiE* gene from *E. coli* genome | TCATTCATCGCCAACTCC |
| 15. | *Ec_thiE*_PCR2_FP | Forward primer to clone *E. coli thiE* gene into pProEX-Hta plasmid | TTCAGGGCGCCATGGATCCGATGTATCAGCCTGATTTTCC |
| 16. | *Ec_thiE*_PCR2_RP | Reverse primer to clone *E. coli thiE* gene into pProEX-Hta plasmid | TACCGCATGCCTCGAGACTGTCATTCATCGCCAACTCC |

**Table S1.** List of primers used in this study.
